# Epigenetic–metabolic axis in the temporal scaling of mammalian cortical neurogenesis across species

**DOI:** 10.1101/2025.09.23.677958

**Authors:** Quan Wu, Charlotte Manser, Taeko Suetsugu, Ryo Yoshida, Hideya Sakaguchi, Yoichi Nabeshima, Hiroshi Kiyonari, Ruben Perez-Carrasco, Fumio Matsuzaki

**Affiliations:** Department of Aging Science and Medicine, Graduate School of Medicine, Kyoto University; Laboratory for Animal Resources and Genetic Engineering, RIKEN Center for Biosystems Dynamics Research, Kobe, Hyogo 650-0047, Japan; Department of Life Sciences, Imperial College London, London, UK; Mathematics Department, King’s College London, London, UK; Neural Organogenesis Laboratory, BDR-Otsuka Pharmaceutical Collaboration Center, RIKEN Center for Biosystems Dynamics Research, Kobe, Hyogo 650-0047, Japan; Division of Experimental Pharmacology, Department of Pharmacology, National Center for Child Health and Development, Tokyo, Japan

**Author notes:** present addresses (Charlotte Manser). present addresses (Hideya Sakaguchi).

## Abstract

Developmental timescales vary widely across species, with mammalian cortical neurogenesis ranging from just days to several months. Given the conserved laminar architecture and regulatory gene expression sequences, the underlying molecular mechanisms controlling neurogenesis rate remain unknown. Using mouse, ferret, and human models, we combined comparative transcriptomics with mathematical modelling to identify conditions that scale temporal gene expression programs during neurogenesis. We show that H3K27me3-mediated repression is critical for maintaining species-specific neurogenesis timescales, and its ablation scales down neurogenesis duration. Furthermore, we identified the tricarboxylic-acid (TCA)-cycle metabolite α-ketoglutarate as a modulator of developmental timing via H3K27me3 demethylation. Together, our findings link epigenetic and metabolic states to the control of species-specific developmental timescales and, ultimately, brain size.

## Main Text

Developmental duration follows the scaling principle, whereby larger species generally require a longer developmental period. Previous studies have shown that differences in the protein synthesis, degradation, and metabolic rates partially explain interspecies variations in processes such as somitogenesis and spinal cord development (*1–6*) However, these processes typically exhibit only a two- to three-fold difference between mice and humans, far less than the order-of-magnitude variation observed in brain development. For instance, the mammalian cerebral cortex develops over 1 week in mice, 1 month in ferrets, and several months in humans (*7–11*). The molecular basis for this temporal scaling remains unclear, although the mechanisms that determine the timing of phase shifts have been extensively studied (*12–16*).

Cortical development in mammals is initiated by the proliferation of neural stem cells (NSCs) in the ventricular zone, followed by neurogenic and gliogenic phases that proceed in a conserved temporal order: deep-layer neurons, upper-layer neurons, and astrocytes (*17–20*). This temporal order is driven by the sequential expression of genes critical for the fate determination of NSCs, many of which are evolutionarily conserved among mammals, including mice, ferrets, and humans (*12*, *21–25*). We previously showed that the temporal pattern of single-cell transcriptomic clusters of cortical progenitors and neurons follows the same conserved sequence between humans (*26*) and ferrets (*22*). To investigate the underlying mechanisms for this scaling, we first performed comparative single-cell transcriptomic analysis over the neurogenic period in all three species and then conducted mathematical modeling to identify the mechanistic conditions that permit such a large variation in neurogenesis timescales.

### Scaling of gene expression pattern with neurogenic duration

We first chose equivalent stages from the mouse, ferret, and human cortices (Materials and Methods, and fig. S1) and calculated developmental pseudotime scores (‘Birthday score’) for each cell of cortical progenitor populations (PAX6+ SOX2+ EOMES-) from single-cell RNA-seq (scRNA-seq) data using previously established temporal gene weights (*12*). The computation of the pseudotimes of cell populations enabled us to mutually compare the biological developmental times among different species. This approach enabled a direct comparison of the biological developmental timing across species. Stage-dependent NSC gene expression patterns aligned closely along the pseudotime, indicating that sequential gene expression in NSCs scales robustly between species (Fig. 1A to C; fig. S2). These findings suggest that the underlying gene regulatory networks (GRN) governing cortical lamination allow scaling along the temporal axis.

**Fig. 1.**
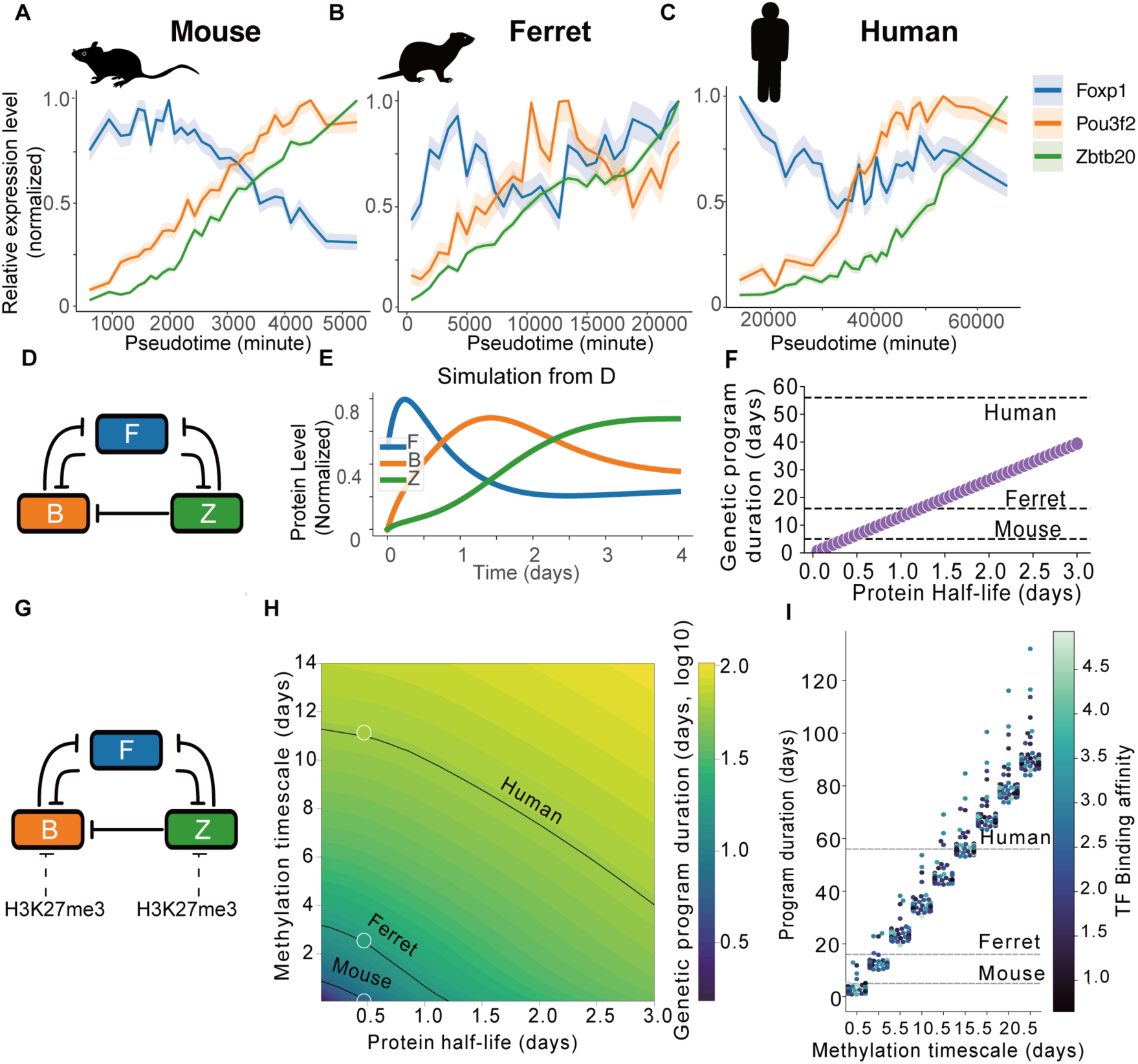
Mathematical models reveal that epigenetically regulated chromatin accessibility allows for the scale of temporal fate transitions across species. **(A to C)** Pseudotime plots of three temporal fate determinants expressed in NSCs from single-cell RNA-seq: Foxp1 (deep-layer neuronal fate), Brn2 (upper-layer neuronal fate), and Zbtb20 (glial fate) in mice (A), ferrets (B), and humans (C). Gene expression levels were normalized and aligned along the developmental pseudotime. (**D-E**) Trajectories of gene expression (E) predicted by the mathematical model given by the corticogenesis GRN Model (D). (F) Prediction of the effect of protein turnover on corticogenesis duration, measured as the time to activate Zbtb20 (fig. S3B). Horizontal dashed lines indicate corticogenesis duration in the three species. (**G and H**) Predicted corticogenesis time (H) for the gene regulatory network (GRN) including histone methylation (G), in which promoter accessibility evolves over time owing to histone methylation and demethylation dynamics (e.g., H3K27me3) for different combinations of protein decay and methylation timescales. Black lines mark parameter combinations matching observed mouse, ferret, and human corticogenesis times (filled circles indicate selected parameters with trajectories shown in fig. S3C). (**I**) Corticogenesis duration as a function of methylation timescale for different transcriptional factor-binding affinity combinations that preserve corticogenesis gene expression sequence. Dashed lines denote the empirical mouse, ferret, and human bands. The parameters and details of the model are provided in the Supplementary Text.

### Protein degradation rates cannot explain differences in tempo (developmental rate)

To investigate the molecular mechanisms underlying corticogenesis at extremely different time scales, we constructed a mathematical model of the cortical gene regulatory network that recapitulates the sequential expression of ‘temporal fate genes’ in NSCs. The model groups genes into three nodes, each identified by a representative gene: Foxp1, Brn2, and Zbtb20, which are critical for producing deep-layer neurons (*27*), upper-layer neurons (*28*), and glial cells (*29*), respectively (Fig. 1D). Their regulatory interactions were inferred by integrating published *in vivo* studies with our data, showing that FOXP1 overexpression suppressed BRN2 and ZBTB20 expression (fig. S3A). The resulting model recapitulates how gene expression levels change over time due to dynamical changes of transcription factors in the network (Fig. 1E).

While the biochemical parameters of the network can modulate developmental tempo, protein degradation is distinctive since it appears linearly in the rate of change of gene expression (Supplementary Text), implying that changes in protein half-life should scale the entire corticogenesis dynamics proportionally(*30*) (Fig 1F). Given that human corticogenesis is over ten-fold slower than that in mice, this would require the human protein half-life to be extended ten-fold. However, the measured protein degradation rates have not demonstrated the capacity to achieve such a disparity(*31*). Thus, while protein turnover can influence the tempo, it cannot account for the dramatic interspecies tempo variation in cortical development.

### Species-dependent rate of H3K27me3 Demethylation during Cortical Development

Protein turnover alone cannot account for the dramatic interspecies differences in developmental tempo, prompting us to consider slower processes as potential regulators. Epigenetic regulation, particularly histone methylation, is an attractive candidate given its inherently longer kinetics. In particular, trimethylation of histone H3 at lysine 27 (H3K27me3), a repressive mark catalyzed by the Polycomb Repressive Complex 2 (PRC2), plays a critical role in regulating the fate transitions of NSCs by suppressing the expression of late-onset genes, which are more highly expressed in E14 than in E11 NSCs, in the early phase of mouse neurogenesis, and PRC2 disruption leads to their premature activation, thereby accelerating the timing of fate transition (19–21, 13).

To test whether methylation dynamics could contribute to tempo scaling, we extended our model to include a dynamic promoter state that decreases accessibility with H3K27me3 accumulation (Fig. 1G). Simulations revealed that both protein degradation and methylation turnover can tune the tempo, but only methylation can do so at realistic biochemical rates. While unrealistically long protein half-lives (2–3 days) would be required to match human tempos, adjusting methylation turnover alone robustly spanned the observed range across the three animal species studied (Fig. 1H, fig. S3C). To ensure that our conclusions were not dependent on a particular parameterization of transcription factor (TF) interactions, we swept the parameters of TF binding affinities and methylation turnover rate, preserving the experimentally observed sequential gene expression order. Within this broad parameter space, binding affinities modulated developmental tempo to a limited extent, but the methylation turnover rate consistently emerged as the dominant determinant (Fig. 1I). These results demonstrate that methylation dynamics provide a parameter-independent modular axis of tempo control, making them a strong candidate mechanism for the large-scale temporal scaling of cortical neurogenesis.

To experimentally explore how histone methylation contributes to the species-specific timescale of neurogenesis, we isolated NSCs from mouse, ferret, and human ESC-derived cortical organoids at matched developmental stages and performed chromatin immunoprecipitation sequencing (ChIP-seq) for H3K27me3. To quantify the (de)methylation dynamics of H3K37me3 during neurogenesis, we identified genomic loci that exhibited dynamic H3K27me3 peak (loss) gain and calculated the average signal change over time for each species. This analysis revealed a markedly faster decline in H3K27me3 levels in mice than in ferrets and humans, consistent with a species-dependent rate of epigenetic changes (Fig. 2, A to E and fig S4).

**Fig. 2.**
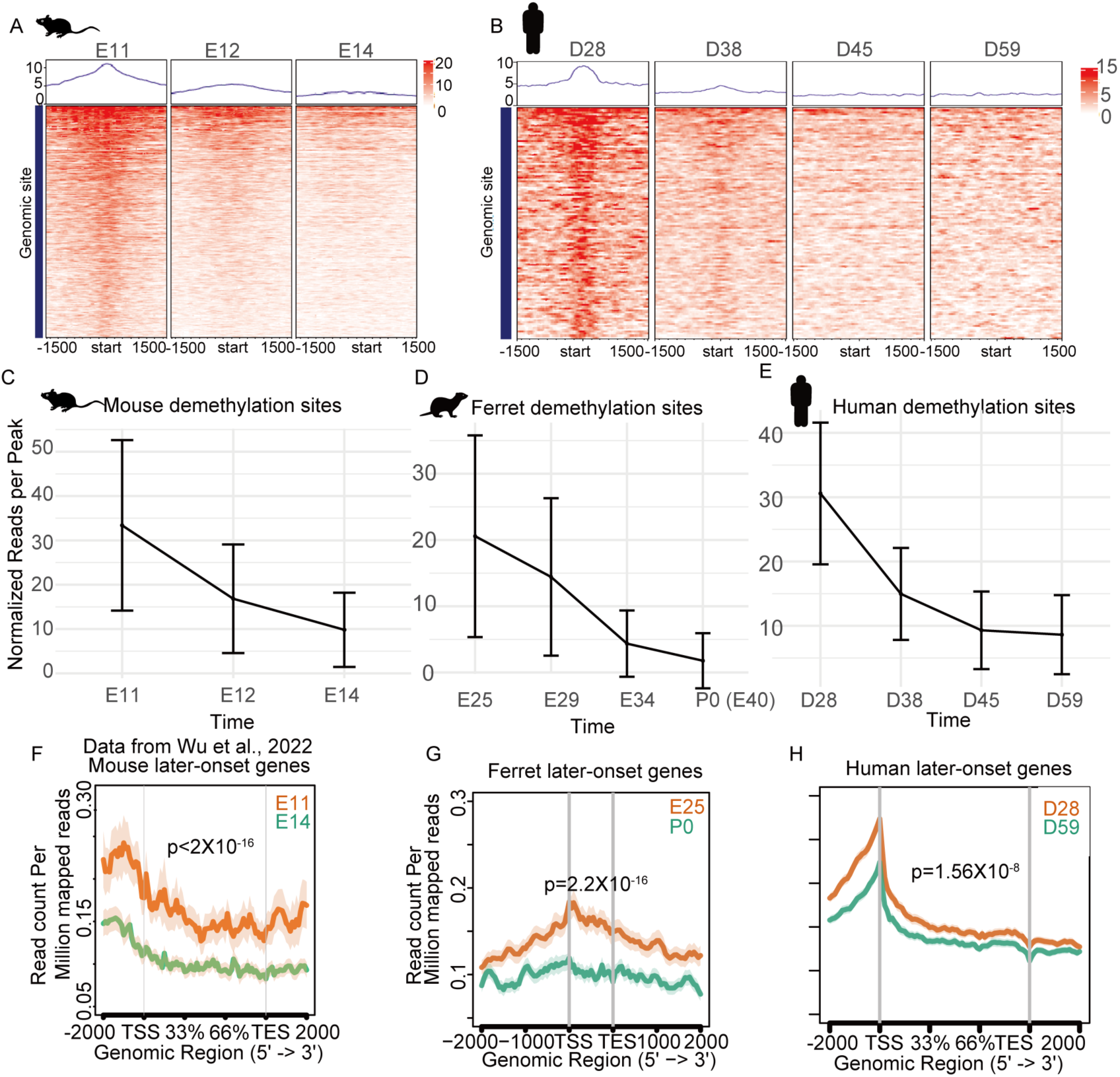
Species-specific demethylation kinetics of H3K27me3 during cortical development. **(A to B)** Heatmap showing the gradual demethylation of H3K27me3 signals over time during the neurogenesis period in mice (A) and humans (B). H3K27me3 signals around the transcription start site with ±1.5 kb flanks were aligned in order of the signal intensity. The color scale indicates the normalized-signal intensity. (**C and E**) Mean normalized reads per H3K27me3 peak ((DiffBind; library-normalized reads/peak) at demethylation sites over the neurogenic time course in mice (C), ferrets (D), and humans (E). Points show the mean across all demethylation sites and replicates; error bars denote ±SD across peaks. Biological replicates used per time point: Mouse: n=3 at E11, n=2 at E12, n=2 at E14.Ferret: n=2 at E25, n=2 at E29, n=3 at E34, n=2 at P0. Human: n=2 at D28, n=2 at D38, n=2 at D45, n=3 at D59. **(F to H**) Meta-profiles of late-onset genes from the transcription start site (TSS) to the transcription end site (TES), scaled to gene length with ±2 kb flanks. The y-axis represents H3K27me3 peak reads per million mapped reads; ribbons indicate ±SD across genes/peaks. The right subpanel shows the mouse data from our previous study (*32*) for comparison purposes. The Wilcoxon signed-rank test was used to evaluate the differences.

Next, we asked whether H3K27me3 dynamics were associated with the activation of late-onset genes by plotting H3K27me3 profiles on late-onset gene sets (defined from the NSC transcriptomes of each species; Table S1). Late-onset genes in mouse, ferret, and human NSCs were enriched for H3K27me3 at early stages and underwent progressive demethylation during neurogenesis, consistent with previous findings in mice (*32*) (Fig. 2, F to H).

### Neurogenic duration scales down by cortical *Ezh2* deletion in mice

To further test the prediction by the mathematical model, we examined whether the depletion of H3K27me3 accelerates cortical neurogenesis, leading to a scale-down of *in vivo* neurogenesis duration. We conditionally deleted the H3K27me3 methyltransferase, Ezh2 (enhancer of zest 2), in the dorsal telencephalon using *Emx1-Cre* (recombination from ∼E9.5). Previous studies have shown that perturbing H3K27me3 methylation promotes the transition from deep- to upper-layer neurogenesis and triggers premature gliogenesis (*12–15*). However, it is unclear whether the duration of the overall neurogenic program is scaled down. To test this as a possibility, we pulse-labeled NSCs with EdU ( or BrdU) once at E12.5, E13.5, E14.5, and E15.5 in the wild-type and *Ezh2* mutants and assessed fate output at E18 (after birth, fewer than 20% of the expected number of *Ezh2*-mutant pups were recovered) to infer the relationship between the birth date and fate output of cortical neurons and glial cells (Fig. 3A). In controls and heterozygous mice, NSCs sequentially generated Tbr1⁺ subplate and layer VI neurons and then Ctip2⁺ layer V neurons from E12.5 to E13.5, followed by layer IV neurons at E13.5 (between Ctip2+ Layer V and Brin2+Layer II/III), and then, predominately Brn2⁺ neurons of layer II–III at E14, with a peak at E15 (Fig. 3, B, D and E; fig. S5A). In contrast, in *Ezh2* mutants, although neuronal stratification was somewhat disordered, neurogenesis was accelerated during E12–E15. At E13, we observed a dramatic reduction in the production of layer IV neurons and an increase in Brn2⁺ (layer II–III) neuron production from NSCs, resembling the output of E14 controls. Brn2⁺ (layer II–III) neuron production peaked at E14, and neurogenesis greatly diminished at E15, mirroring the control E15 and E16, respectively (Fig.3, B, D, and E).

**Fig. 3.**
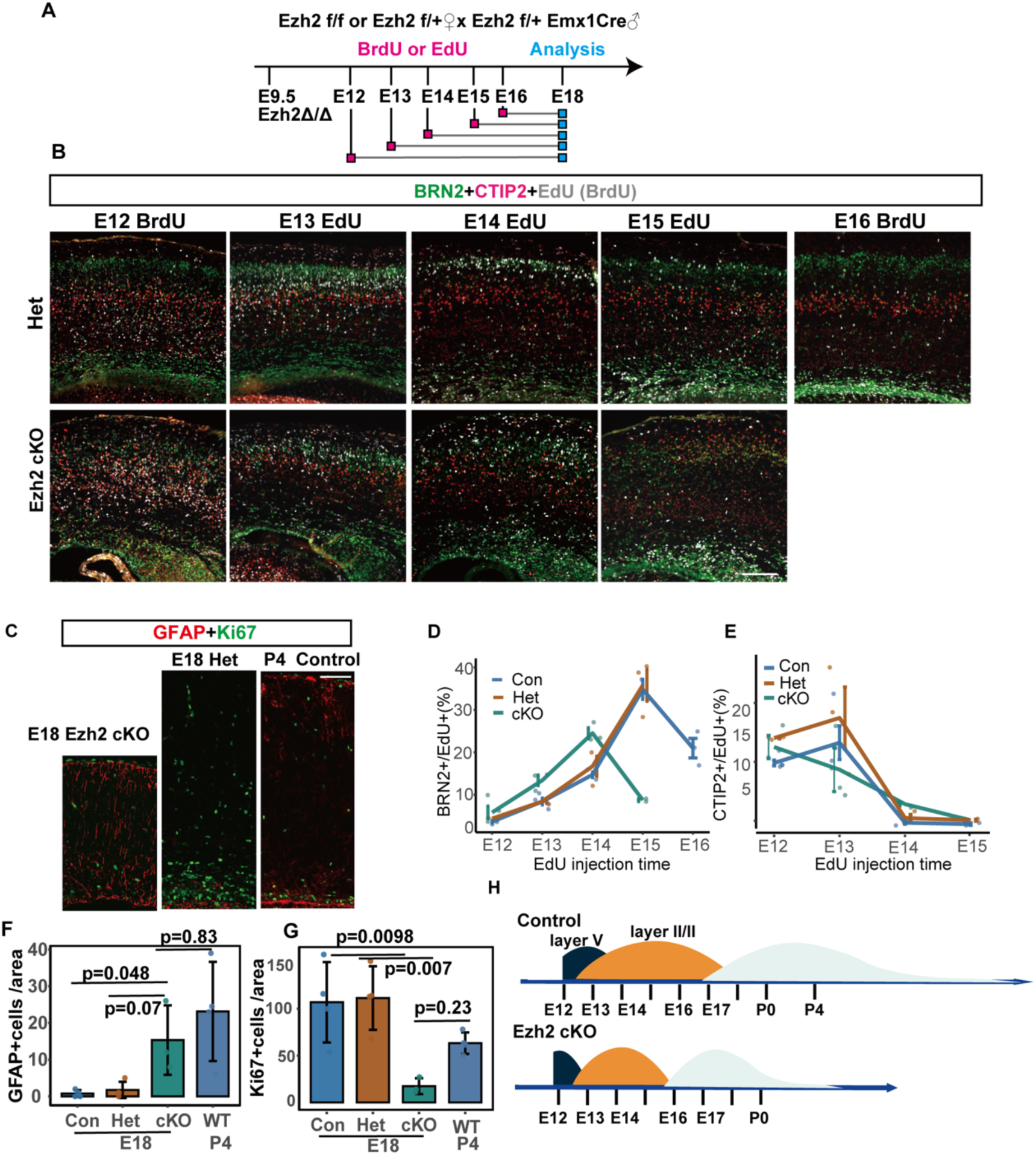
Loss of Ezh2 scales neurogenesis and gliogenesis in the embryonic mouse cortex. **(A)** Experimental timeline of this study. *Ezh2* conditional knockout (cKO) mice and control littermates were administered a single EdU (BrdU) pulse at E12.5 (BrdU), E13.5, E14.5, E15.5, and E16.5 (control only), and the brains were analyzed at E18.5. **(B)** Representative coronal sections showing triple staining for BRN2, CTIP2, and EdU/BrdU (birth-dating) after pulses on the indicated days. Scale bar, 50 µm **(C)** Immunostaining for GFAP (astroglia) and KI67 (mitotic) at E18 in heterozygous and *Ezh2* cKO mice and at P4 in the control. Scale bar, 50 µm. **(D and E)** Quantification of the percentage of BRN2^+^EdU^+^ cells (D) and CTIP2^+^EDU^+^ cells (E) among labelled neurons at the indicated pulse times in control, heterozygous, and *Ezh2* cKO mice. Bars show mean ± SEM. Data points denote the biological replicates. Sample numbers: (Con): n=3 at E12, n=4 at E13, n=3 at E14, n=2 at E15, n=3 at E16; (Het): n=3 at E12, n=3 at E13, n=3 at E14, n=3 at E15; (cKO): n=4 at E12, n=3 at E13, n=3 at E14, n=2 at E15. **(F–G)** Quantification of GFAP and KI67 cell numbers per area at the late embryonic/early postnatal stages. Bars represent the mean ± SD, and points represent biological replicates. Pairwise comparisons of GFAP used ANOVA tests with Tukey HSD correction; Pairwise comparisons of KI67 used Kruskal–Wallis tests with Dunn’s test with Holm correction. The adjusted p-values are indicated. Con: n=4; Het: n=4; cKO: n=3. **(H)** Schematic summary of the scaling down of neurogenic and gliogenic programs by the knockout of *Ezh2*.

At E18, Ezh2-deficient cortices showed a marked reduction in KI67 ⁺ proliferating cells and TBR2 ⁺ intermediate progenitors (Fig. 3C and fig. S5, B and C), suggesting premature termination of neurogenesis. In addition, we observed robust GFAP upregulation at E18, comparable to that in control P4, indicating an advanced onset of gliogenesis (*33*), and consequently shortening the time window of neurogenesis (Fig.3, F and G, and fig. S5B). Consistently, brain size was significantly reduced in the mutants (fig. S5, D and E). Together, these Ezh2 phenotypes strongly support a temporal scale-down of cortical neurogenesis upon a reduction of H3K27me3 methylation in mouse corticogenesis (Fig.3 H)

### Reducing H3K27me3-methylation increases the rate of cortical neurogenesis in ferrets

Next, we examined whether H3K27me3 affects cortical neurogenesis progression in ferrets. We introduced vectors for CRISPR/Cas9-dependent knockout of the *EZH2* gene, as well as an EGFP vector, via *in utero* electroporation in ferrets at approximately E30–E32, the timing of deep-layer neurogenesis, and confirmed substantial reductions in both EZH2 protein and H3K27me3 levels (fig. S6). Analysis of EGFP^+^ cells using single-cell RNA sequencing at E34 and E40 (Fig. 4A), revealed that TLE4*^+^*layer VI neurons and mature upper layer neurons (CUX2 high expression) were generated in the wild type. In contrast, EZH2-depleted cells showed a significant reduction in TLE4⁺ deep-layer neurons and an increase in CUX2 high-expressed upper-layer neurons, suggesting an accelerated neurogenic progression (Fig. 4B-F and fig. S7A-D). In alignment with these findings, immunostaining also suggested a reduction in TBR1^+^ deep-layer neurons and a trend toward an increase in SATB2^+^ upper-layer (fig. S7 E to G). Moreover, progenitors transduced with either gRNA set exhibited higher birthday scores than controls, with the increase more pronounced for gRNA set 1, consistent with its greater Ezh2-knockout efficiency (Fig. 4G). Genes associated with astrogenesis, SPARCL1 and LGALS1, were upregulated in Ezh2-deficient NSCs as early as E40, whereas their expression was weak in wild-type cells at E40 (Fig. 4J and K). Furthermore, at P10, GFAP⁺ astrocytes were slightly increased in EZH2 KO cells compared to controls. These observations are consistent with the earlier occurrence of gliogenesis in Ezh2 mutant cells than in wild-type cells (Fig. 4H and I,). Overall, our data suggest that the loss of Ezh2 shifts both neurogenic and gliogenic programs earlier in ferret corticogenesis, consistent with the phenotypes observed in *Ezh2* mutant mice.

**Fig. 4.**
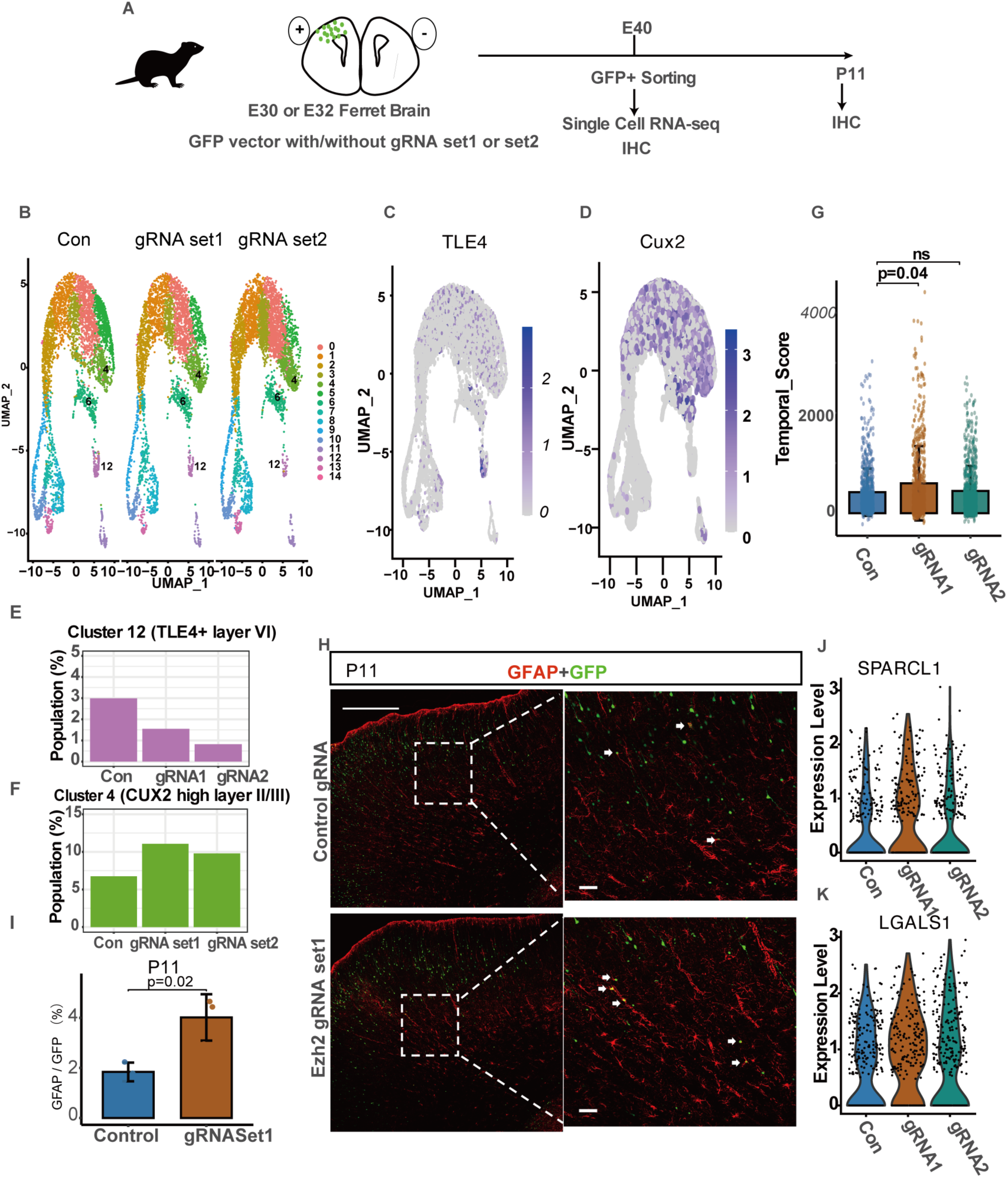
Loss of Ezh2 accelerates temporal fate progression in the developing ferret cortex. **(A)** Experimental design of this study. E30/E32 ferret embryos were electroporated with expression vectors for GFP, *Ezh2*-gRNA set 1 or set2, and the CRISPR/Cas9 system. **(B)** UMAP projection of single-cell RNA-seq data from ferret cortices at E40 following *in utero* electroporation (IUE) at E32 with no gRNA (CON) or two independent Ezh2-targeting gRNA sets (gRNA1 and gRNA2). (**C and D)** Normalized expression of TLE4 (layer VI) and CUX2 (layer II/III) across the UMAP. **(E and F)** Fraction of cells assigned to the TLE4⁺ (cluster 12) or CUX2-high (cluster 4) clusters for each condition. **(G)** Plots of temporal scores for each condition (dots indicate individual cells). Pairwise comparisons used two-sided Wilcoxon rank-sum tests with Benjamini–Hochberg FDR correction; the adjusted p-values are indicated. **(H)** Representative coronal sections of ferrets stained for GFAP (red) and electroporated GFP (green) at P11. Scale bar, 500 µm (left panels). Scale bar, 50 µm (right panels). Arrows indicate GFAP^+^ cells. Student’s t-test was used to calculate p-values. n=3 for each condition. **(I)** Quantification of the GFAP⁺ fraction in GFP cells at P11. Bars indicate mean ±SD. **(J, K)** Violin plots of SPARCL1 and LGALS1 expression for each condition at E40.

### α-ketoglutarate (α-KG) regulates H3K27me3 levels of temporally changing genes during neurogenesis across species

Next, we sought to identify the factors influencing the H3K27me3 demethylation rate in different species. Demethylation of H3K27me3 is catalyzed by Fe(II)/α-ketoglutarate-dependent Jumonji-type (JmjC) dioxygenases (*34–36*). Previous studies have shown that intracellular α-ketoglutarate (α-KG) levels modulate the global methylation status in embryonic stem (ES) cells, induced pluripotent stem (iPS) cells, and cancer cells (*37–40*), raising the possibility that α-KG concentration regulates the rate of H3K27me3 demethylation in the developing cortex. To test this, we first cultured NSCs from the developing mouse cortex for three days with exogenous α-KG (Fig. 5A), and found that higher α-KG concentrations significantly reduced H3K27me3 levels (Fig. 5B), confirming its ability to promote demethylation in NSCs. α-KG treatment also increased the transcripts for glycolytic enzymes: hexokinase 2 (HK2), enolase 1 (ENO1), and lactate dehydrogenase A (LDHA), and glucose transporters (SLC2A1/GLUT and SLC2A3/GLUT3), consistent with enhanced glycolytic flux (fig. S8 A-C and Table S3).

**Fig. 5.**
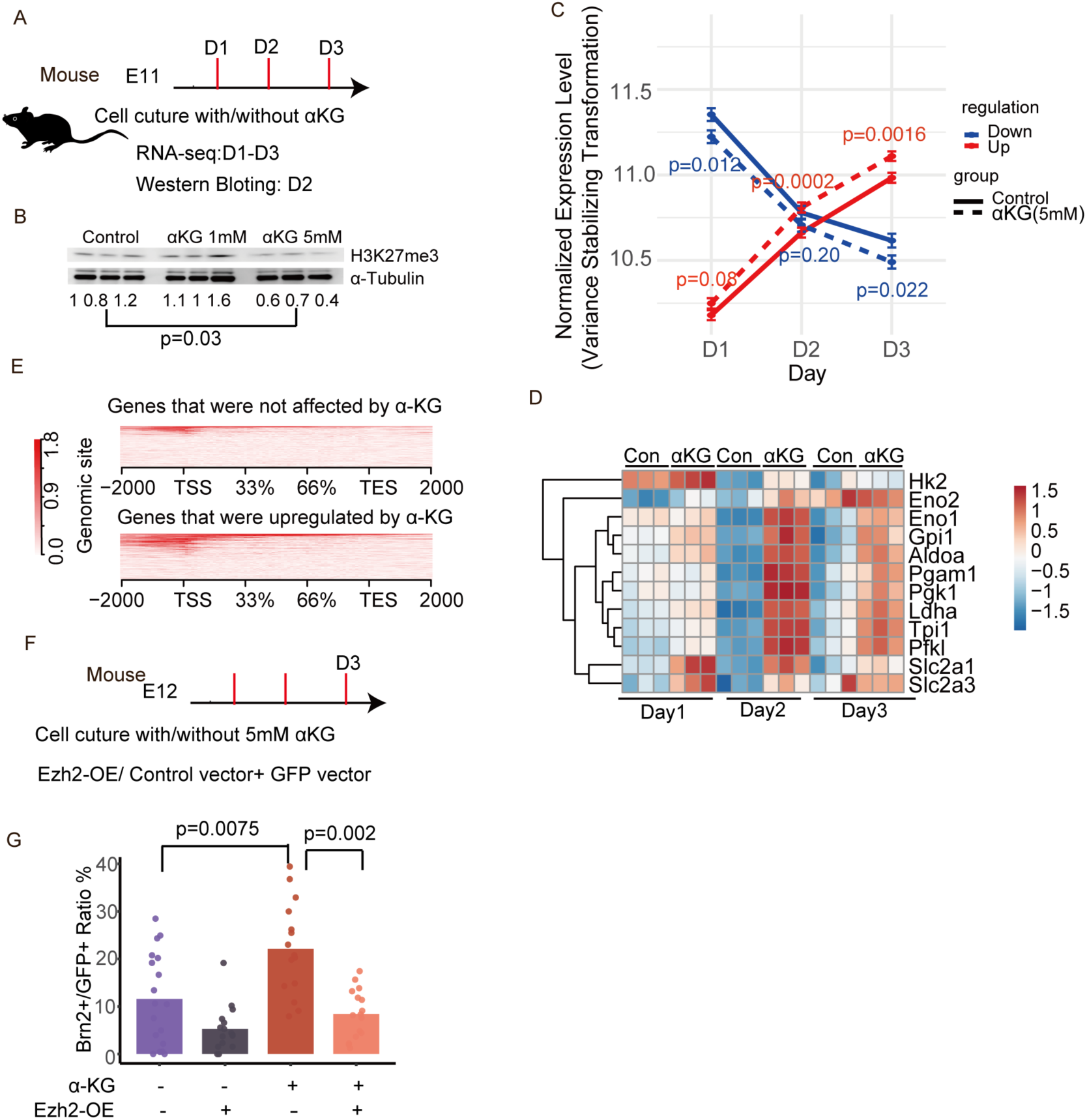
α-Ketoglutarate (α-KG) promotes the demethylation of H3K27me3. (**A**) Experimental design for the treatment of mouse E11 cortical cells with or without α-KG. (**B**) Western blot analysis for H3K27me3 with α-tubulin loading control at D2. The numbers indicate H3K27me3/α-tubulin intensity normalized to that of α-tubulin. Student’s t-test was used to calculate the p-value. (**C**) Mean normalized expression of differentially expressed genes (DEGs) over time in control (solid lines) and 5 mM α-KG (dashed lines) cultures are shown separately for upregulated (red) and downregulated (blue) gene sets. Error bars represent mean ± SEM; Wilcoxon signed-rank test for the indicated day-to-day comparisons. N=3 for each condition in each day. (**D**) Heatmap of representative metabolic/glycolytic genes across D1–D3 period. (**E**) Metagene profiles of H3K27me3 (from E12 NSCs) across gene bodies for genes unaffected by α-KG (top) versus genes upregulated by α-KG (bottom) at D1. (F) Design of rescue assay. (G) Percentage of BRN2⁺ among GFP⁺ cells at D3 under the four conditions. Bars show mean ± SD; points represent counted regions. Group differences were analyzed using the Kruskal–Wallis test, followed by pairwise Dunn’s tests with Benjamini–Hochberg FDR correction.

### α-KG regulates temporal shifts in gene expression during neurogenesis

Next, we examined whether increasing α-KG concentration by exogenous α-KG addition accelerates shifts in gene expression during neurogenesis using cultured mouse NSCs. We found that genes that were upregulated from D1 to D3 were further upregulated by α-KG, whereas downregulated genes were further repressed (Fig. 5C and D), indicating that α-KG accelerates the gene expression during neurogenesis *in vitro*.

To evaluate whether α-KG has similar effects also in species with a slower developmental rate, we cultured ferret embryonic cortical cells from E32 (deep-layer neurogenesis stage) for two and six days with or without α-KG, followed by RNA sequencing (fig. S9A). α-KG treatment induced widespread transcriptional changes, including significant downregulation of early-onset genes and upregulation of late-onset genes, consistent with an overall acceleration of developmental timing in ferret cortical cells *in vitro* (fig. S9B and Table S4).

As α-ketoglutarate (α-KG) is a cofactor for demethylases, including the H3K27-specific KDM6 family and DNA-demethylating TET enzymes, we tested whether α-KG modulates neurogenic gene expression by promoting H3K27me3 demethylation. This predicts that the genes upregulated by α-KG treatment should be H3K27me3-enriched at the corresponding developmental stage. Consistent with this, in both mice and ferrets, genes induced by α-KG carried significantly higher H3K27me3 levels than genes whose expression did not change (Fig 5E, fig. S8D and fig. S9C).

To directly test whether α-KG is involved in determining the timing of phase shifts in neurogenesis through H3K27me3 demethylation, we cultured E12 mouse cortical cells for three days with α-KG treatment, with or without the overexpression of the methyltransferase Ezh2. While α-KG treatment led to a marked increase in Brn2^+^ cells, overexpression of Ezh2 largely suppressed this upregulation, indicating a compensation of α-KG effect on Brn2 expression by an increase in Ezh2 levels (Fig.5 F and G). These results provide strong evidence that α-KG promotes late-onset genes during the later neurogenic phase, at least partially through H3K27me3 demethylation.

### Comparison of temporal control in neurogenesis between mouse and human

If α-KG and EZH2 function operate in human cortical development as in mice, α-KG and EZH2 inhibitor GSK343 must show a similar effect on neurogenesis tempo. We tested this hypothesis using cortical organoids developed from human ESCs. When GSK343 or α-KG was applied to cortical organoids from 34 days (D34) of culture, both α-KG- and GSK343-treated organoids exhibited accelerated production of upper-layer neurons by D52 (Fig.6, A and C) and GFAP astrocytes by D65 (Fig. 6, B and D). This result supports the conserved role of α-KG-dependent H3K27me3 demethylation in controlling the timing of phase shifts during neurogenesis and the onset of gliogenesis across species, leading to neurogenic scaling.

**Fig. 6.**
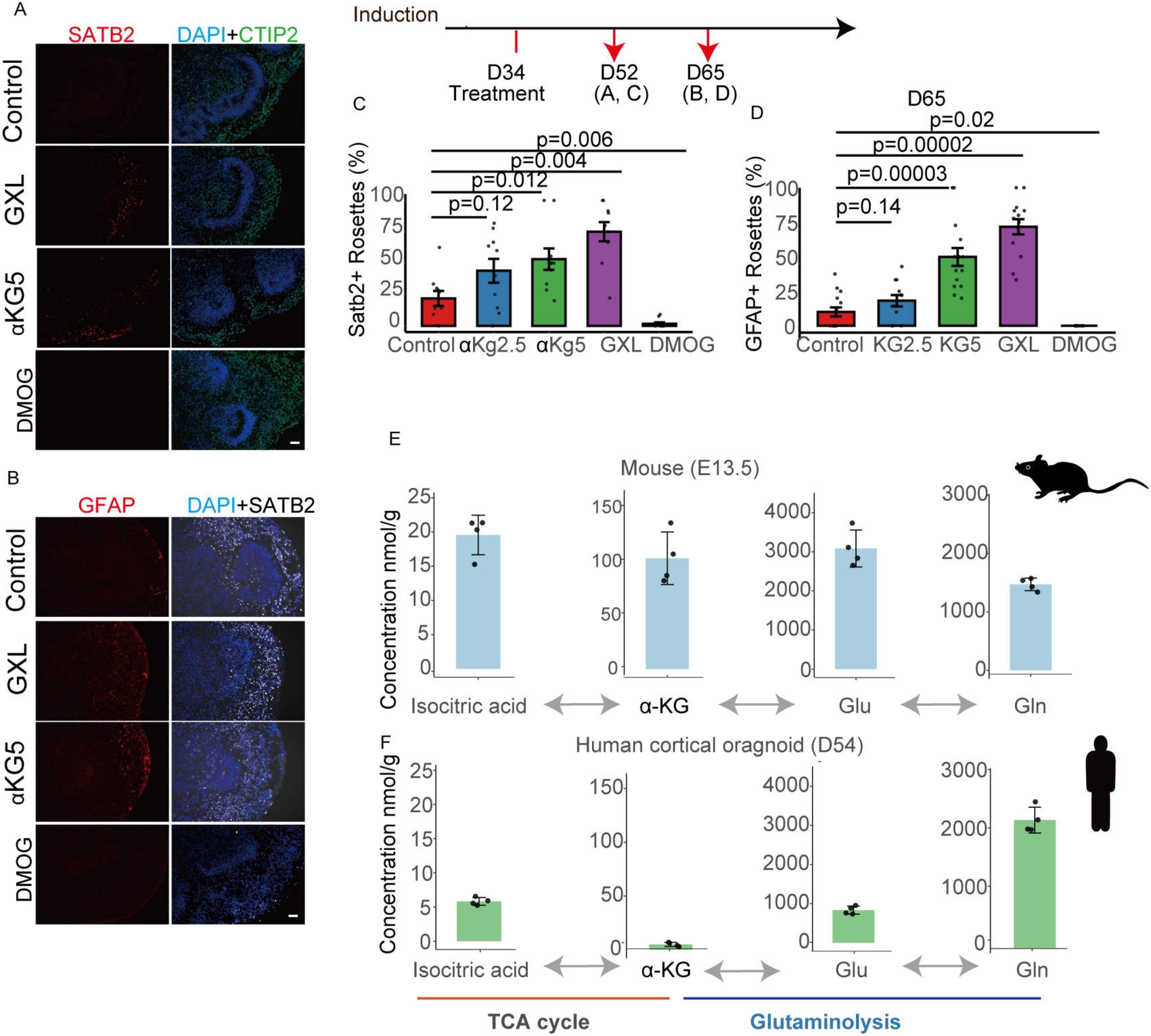
α-KG advances upper-layer neurogenesis and gliogenesis in human cortical organoids while DMOG suppresses it. **(A and B)** Representative immunofluorescence images of SATB2/CTIP2/DAPI (A) and GFAP/SATB2/DAPI (B) in the treatment groups. Scale bar, 50 µm. **(C-D)** Quantification of SATB2⁺ rosettes (% of total rosettes) and GFAP⁺ rosettes (%) in organoids under the indicated conditions. Bars represent the mean ± SD, and the points represent different organoids from two independent inductions. Group differences were assessed using the Wilcoxon rank-sum test, followed by pairwise Dunn tests with Benjamini–Hochberg FDR correction. **(E–F)** Targeted metabolomics of human organoid (D54) and mouse cortical (E13) samples: concentrations (nmol/g) of α-KG and related metabolites along the TCA cycle (isocitric acid) and glutaminolysis (Gln and Glu).

We confirmed the effect of α-KG on the late-onset gene induction by using α-KG antagonist Dimethyloxallyl Glycine (DMOG) in both mouse cortical NSCs and human cerebral organoids. E11 mouse cortical cells treated with DMOG showed increased H3K27me3 levels and suppressed induction of late-onset genes, such as *Zbtb20* and *Pou3f2*, while maintaining the expression of early-onset genes, such as *Hmga2* and *Dmrt3* (fig. S10, A to E). We also confirmed that DMOG suppressed the fate transition from layer V neurons to layer IV neurons in mice (fig. S10, F-I). In human cortical organoids, DMOG treatment dramatically inhibited the emergence of upper-layer neurons and glial cells, as observed in mice (Fig.6, A‒D). Together, these results suggest that α-KG–dependent demethylation of H3K27me3 is required for the progression of neurogenic programs across species with widely different developmental timescales.

### Different Metabolic Rate across species

To investigate whether differences in α-ketoglutarate (α-KG) metabolism contribute to the slower rate of human neurodevelopment, we compared metabolomic profiling between E13 mouse cortices and D54 human cortical organoids. The absolute concentration of α-KG in human organoids was dramatically lower than that in developing mouse brains; in fact, it was almost undetectable in human samples (Fig 6E). Other TCA cycle intermediates were also consistently reduced in human cortical organoids compared to developing mouse cortical tissues, suggesting a global reduction in mitochondrial flux in humans compared to mice (fig. S11 and Table S5). The overall low activity of the TCA cycle in human organoids may be relevant to the much slower developmental rate of humans, while comparisons between cortical organoids and *in vivo* mouse brains do not exclude the possibility of culture-specific effects. As for glutaminolysis that also produces α-KG, the upstream substrates (glutamine and glutamate) of this pathway are present at certain levels in contrast to a nearly negligible level of α-KG (Fig 6E), implying inefficient conversion of glutamate to α-KG in human cortical organoids. Consistent with this, the expression of GLUD1, the enzyme catalyzing glutamate-to-α-KG conversion, was markedly lower in human NSCs than in mouse and ferret NSCs, as revealed by single-cell RNA-seq data (fig. S12).

## Discussion

In this study, we revealed conserved mechanisms that scale neurogenesis duration—temporal scaling—across species from mice to humans spanning over a 10-fold difference. Mammals share many developmental programs, including corticogenesis, despite vast differences in developmental timescales between species. However, it remains unclear what controls the duration of neurogenesis and gliogenesis in a scalable manner (preserving the 6-neuronal architecture of the mammalian cerebral cortex). Our results in Ezh2 mutant mice provide the first evidence that H3K27m3 methylation-demethylation (catalyzed by EZH2 and KDM6, respectively) plays a key role in this process. Ezh2 knockout not only accelerates the timing of the neuro-gliogenesis transition but also shortens the neurogenesis duration, as judged by an approximately one-day shift in neurogenic phases and four-days premature gliogenesis initiation. We further demonstrated that EZH2 knockout accelerates neurogenesis in ferrets, suggesting that the role of EZH2 is conserved across mammals. We were unable to determine whether gliogenesis was also shortened because most Ezh2 mutant mice died soon after birth.

Our mathematical model including epigenetic modification, indicates that such mechanisms can operate as a general means of establishing developmental programs that are scalable among species with a wide range of timescales. This is true for mammalian cortical development, as shown in this study. This has not been found in processes where timing differences between species are relatively small, such as in somite formation or spinal cord development(*3*, *4*). Thus, our combination of mathematical modeling and experimental evidence supports the role of epigenetic dynamics in temporal scaling for cortical neurogenesis, providing a fascinating evolutionary viewpoint on epigenetic mechanisms.

Our study also revealed that α-KG levels contribute to shortened neurogenesis by enhancing the demethylation of H3K27me3 as a cofactor of KDM6, a JmjC-type dioxygenase. This type of enzymes functions in catalyzing numerous epigenetic processes beyond the removal of H3K27me3, including the demethylation of H3K4me3, H3K9me3, and DNA methylation. Additionally, other metabolic cofactors, such as S-adenosylmethionine (SAM), a methyl donor, and acetyl-CoA, a substrate for histone acetylation (e.g., H3K27ac), play critical roles in influencing epigenetic dynamics(*41–43*). These observations suggest the possibility that developmental programs of an extended duration, such as those in humans, may arise from a combination of slower α-KG–dependent epigenetic demethylation reactions and metabolic activities, thereby affecting multiple epigenetic and gene regulation systems (*44–47*).

## Acknowledgments

We thank Z.Q. Yang, Y. Yamaga and all members of the Matsuzaki Laboratory for their discussions and technical advice. We thank the Single-cell Genome Information Analysis Core (SignAC) at WPI-ASHBi, Kyoto University, for their support. We thank C. Tanegashima, K. Tatsumi, and O. Nishimura of the Laboratory for Phyloinformatics, RIKEN Center for Biosystems Dynamics Research (BDR) for NGS library preparation, sequencing, and assistance with data production and analysis. We thank A. Sehara for her critical suggestions regarding the data analysis. We thank H. Miyachi and S. Kitano for helping with the IVF of mutant mice.

## Author contributions

Conceptualization: F.M. and Q.W.; Data curation: F.M. and Q.W.; Formal analysis: Q.W., T.S., and C.M.; Funding acquisition: F.M., H.S. and Q.W.; Investigation: Q.W., R.Y, H.S. and T.S.; Methodology: R.P.C., Q.W. and C.M. Project administration: F.M. and Q.W.; Resources: H.K, R.Y and H.S. Software: Q.W. and C.M; Supervision: F.M and R.P.C; Validation Visualization: Q.W. and C.M.; Writing – original draft: F.M., Q.W and C.M; Writing – review & editing: F.M., Q.W., R.P.C, and C.M.

## Funding

Japan Society for the Promotion of Science (JSPS) Grants-in-Aid for Scientific Research (KAKENHI) grand H00383 and H04003 (F.M.)

Japan Society for the Promotion of Science (JSPS) Grants-in-Aid for Scientific Research on Innovative Areas (Research in a Proposed Research Area) H04791 (F.M.)

Japan Society for the Promotion of Science (JSPS) Grants-in-Aid for Scientific Research (KAKENHI) grand K05787 (Q.W.)

Japan Agency for Medical Research and Development (AMED) Interstellar Initiative and Interstellar Initiative Beyond (Q.W.)

Takeda Science Foundation (Q.W.)

Japan Science and Technology Agency PRESTO Sakigake program JPMJPR2382 (Q.W.) Japan Society for the Promotion of Science (JSPS) Short Term Pre/Postdoctoral Fellowship (C.M.)

BBSRC grant BB/Y002709/1 (R.P.C.)

Leverhulme Trust RPG-2023-085 (R.P.C.)

RIKEN Center for Biosystems Dynamics Research Organoid Project (Q.W., F.M. and H.S.)

## Data and materials availability

All raw and proceeded data for scRNA, ChIP-seq and RNA-seq are available at the DNA Data Bank of Japan

## Supplementary Materials

### Materials and Methods

#### Animals

All animal procedures were performed in accordance with the guidelines for animal experiments at Kyoto University and RIKEN in Japan. All animal experiments were approved by the Institutional Animal Care and Use Committee of RIKEN Kobe Branch and Animal experiment committee of Kyoto University.

Ezh2 conditional knockout mice and Emx1-Cre mice were reported previously(*1*, *2*) and received from RIKEN BRC (strain name: B6;B6129-Ezh2<TM1YUGO>/Hko and B6.129P2-Emx1<tm1.1(cre)Ito>/ItoRbrc, respectively). These mice were maintained on a C57BL/6 background. Wild-type mice used for the inhibitor treatment were maintained on an ICR background.

Ferrets were purchased from Marshall Bioresources (New York, United States) and SLC-Japan (Hamamatsu, JAPAN). Ferrets (both male and female) kept in a short photoperiod room (8 h of light) were transferred to a long photoperiod room (16 h of light) to induce estrus. After confirming estrus in the females, mating was performed on the next day. The day after mating was designated embryonic day 0 (E0).

#### Cross-species developmental timing inference among mouse, ferret, and human

To examine the conserved and species-specific temporal dynamics of cortical neurogenesis, we established cross-species developmental stage correspondence among mice, ferrets, and humans. For the mouse, we selected embryonic day (E)11 to E16, covering the initial generation of deep-layer (VI) neurons through to the almost end of production of upper-layer (II/III) neurons, as determined by EdU pulse-labeling experiments(*3*) (fig. S1A). For the ferret, we selected embryonic day 25 (E25) to postnatal day 1 (P1), which aligns with mouse E11 to E16, based on BrdU birth-dating analysis (*4*) (fig. S1B).

Given the lack of equivalent day-resolution data for humans, we employed a Seurat–based label transfer approach (*5*) to project developmental stage annotations from human single-cell RNA-seq reference datasets (with known gestational week [GW] metadata) onto ferret cells. Each ferret neural stem cell (NSCs) was assigned a predicted human gestational stage along with a label transfer confidence score, which enabled quantitative interspecies alignment.

For each ferret developmental stage, we computed the distribution of predicted human GW annotations and visualized the proportions as a stage-by-stage heatmap (fig. S1C). For all ferret stages, we calculated the weighted mean human gestational week using confidence scores as weights, producing a trajectory plot of the predicted human developmental timing across the ferret neurogenic timeline (fig. S1D). The resulting heatmap and trajectory plot revealed that the majority of ferret cells from E25 through P1 mapped consistently to human stages between GW10 and GW18. This empirical mapping provides a rational basis for selecting the gestational window as the primary reference frame for comparing ferret and human cortical development.

#### Pseudotime estimation based on temporal fate gene expression across species

To quantify the developmental progression of cortical NSCs across species, we computed a pseudotemporal metric termed “Birthday score” based on previously reported temporal fate genes(*6*). Single-cell RNA sequencing (scRNA-seq) datasets from mice, ferrets, and humans at selected stages were used (see above section).

Raw count matrices were normalized, variable features were selected, and the data were scaled using Seurat’s default pipeline. A curated list of temporal fate genes was obtained from a previous study(*6*), each associated with a numerical weight representing its contribution to developmental progression. Genes that were absent from the dataset were excluded. The final gene sets shared across species were intersected and used for downstream scoring.

For each cell, a weighted sum of the log-normalized expression values of the temporal genes was computed as follows:

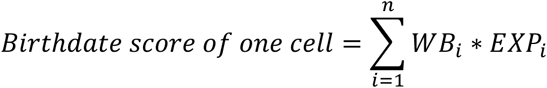

*WB_i_* and *EXP_i_* indicate the weight of each temporal-related gene(*i*) and its expression level in each cell, respectively.

By normalizing each cell’s weighted “Birthday score” within its species-specific score range (setting the minimum to 0 and the maximum to 1) and then mapping it onto the actual developmental time in minutes, we obtained a cross-species comparable pseudotime value that reflects the progression of neurogenesis.

#### Isolation of NSCs from ferret and human cerebral organoids

To purify NSCs populations in ferrets, carboxyfluorescein diacetate succinimidyl ester CFSE ; C34554, Invitrogen) labeling was performed as follows. CFSE dye (final concentration 5 mM) was prepared by dissolving 18 µL of DMSO into the stock CFSE reagent. A working solution was prepared by mixing 1 µL of CFSE with 50 µL of saline and 2.5 µL of 10% First Green.

This labeling mixture was injected into the brains of live ferret embryos. Following injection, the embryos were incubated at room temperature (RT) for 30 min to allow for cellular uptake and ester hydrolysis of the dye. After incubation, the dorsal forebrain regions were dissected from the embryos. For early stage ferret embryos (E25 and E29), tissue dissociation was performed by incubation with 0.25% trypsin in HBSS at 37 °C for 10 min. The cortical plate was carefully removed before dissociation of the later-stage ferret embryos (E34 and P0). These samples were incubated with 130 U of papain (Worthington, Cat# LK003176) in 5 mL of HBSS at 37 °C for 30 min. A neural tissue dissociation kit (Sumitomo Bakelite, MB-X9901) was used to dissociate human cortical organoids. Dissociated NSCs were isolated using a cell sorter based on CFSE and Venus signals (SH800, SONY).

#### Chromatin Immunoprecipitation (ChIP) and Library Preparation

ChIP-seq was performed as previously described (*7*). NSCs were fixed in 0.25% paraformaldehyde for 10 min at room temperature and quenched with 0.1 M glycine. After centrifugation (1,500 × g, 5 min), the cells were resuspended in ChIP buffer (10 mM Tris-HCl pH 8.0, 200 mM KCl, 1 mM CaCl₂, 0.5% NP-40) and briefly sonicated. Chromatin was digested with micrococcal nuclease (50 U/mL, Worthington) at 37 °C for 20 min, and the reaction was stopped with 10 mM EDTA. After centrifugation (15,000 × g, 5 min), the pellets were lysed in RIPA buffer and sonicated. For each ChIP, 100 μL of lysate was incubated with Dynabeads (Invitrogen) pre-bound with 1 μL of anti-H3K27me3 (CST #9733) antibodies. Following overnight incubation at 4 °C, the beads were washed with low- and high-salt buffers, and chromatin was eluted at 65 °C in elution buffer (50 mM Tris-HCl pH 8.0, 10 mM EDTA, 1% SDS) and reverse crosslinked. DNA was treated with RNase A and proteinase K, followed by phenol:chloroform extraction. ChIP-seq libraries were prepared using the KAPA LTP Library Prep Kit with input (2 ng) or full ChIP DNA, and amplified using the KAPA Real-Time Library Amplification Kit to minimize PCR cycles.

Library preparation for ChIP-sequencing (ChIP-seq) was performed using the KAPA LTP Library Preparation Kit (KAPA Biosystems) and 2 ng of input DNA or the entire amount of ChIP DNA obtained was used. KAPA Real-Time Library Amplification Kit (KAPA Biosystems) was used in conjunction with the library preparation kits described above to minimize the number of PCR cycles for library amplification

#### Bioinformatics analysis of ChIP data

Raw sequencing reads were aligned to the human genome (hg19) and ferret genome(*8*) using Bowtie2, and peaks were called using MACS2. Differential binding analysis and count normalization were conducted using the *DiffBind* package (v3.10.1) in R(*9*). Mouse data were obtained from our previous study (*7*).

A sample sheet defining the experimental metadata (time point, replicate, condition) was used to generate a *DBA* object, from which the read counts were computed per peak for each sample. PCA and correlation plots were generated to assess the sample clustering and quality. The dba.contrast and dba.analyze functions were used to define time-series contrasts and identify differentially methylated regions (DMRs) across developmental time. Differential peaks were extracted using dba.report, and the top 1,000 most significant DMRs (based on the false discovery rate (FDR)) were selected for further analysis.

To extract temporal dynamics, normalized read counts for each selected peak were extracted and subjected to polynomial regression analysis using the *maSigPro* package. A design matrix was created from the sample metadata (make.design.matrix), and generalized linear modeling (p.vector followed by T.fit) was used to fit the expression trajectories over time. Significant dynamic peaks were selected based on R² > 0.3 and FDR < 0.05, and hierarchical clustering was performed to identify groups of peaks with similar temporal demethylation patterns. Temporal H3K27me3 coverage across peak clusters was visualized using line plots with error bars, showing the average signal ±SD per time point. For each cluster, peak-wise coverage was computed, reshaped using the melt function, and merged with the sample metadata. The summarized data were plotted using ggplot2.

#### Isolation and meta-profiles of late-onset genes in ferrets and humans

To identify late-onset genes in human cerebral organoids, we isolated NSCs from early (day 24, D24) and late (day 59, D59) organoids and performed bulk RNA-seq. Genes showing >1.5-fold higher expression at D59 than at D24, with an adjusted P-value < 0.05, were designated as late-onset. For ferrets, late-onset NSC genes were defined from scRNA-seq by comparing NSCs at E25 and postnatal day 0 (P0) using the same criteria as above. The resulting gene lists are provided in Table S1.

#### Immunohistology and confocal imaging

Immunohistological analysis was performed as previously described (*7*).Briefly, dissected brains were fixed overnight in 1% paraformaldehyde (PFA), cryoprotected with 25% sucrose, and embedded in OCT compound (Tissue-Tek; Sakura). Tissue sections (12 µm) were cut using a cryostat (Leica Microsystems, Wetzlar, Germany). For immunostaining, the sections were treated with HistoVT One (Nacalai Tesque) at 90°C for 15 min and blocked for 1 h at room temperature in PBST (0.1% Tween20 in PBS) containing 3% skim milk. The samples were then incubated overnight at 4°C with primary antibodies at the appropriate concentrations. The sections were then exposed to fluorescently labeled secondary antibodies (1:400; Alexa Fluor 488, Cy3, or 647; Jackson ImmunoResearch) for 60 min at room temperature. Nuclei were counterstained with DAPI. A scanning confocal microscope (Olympus FV1000 or Nikon A1) and a microscope (Keyence BZ-X) were used for observation. For TBR2/EOMES and KI67 staining, the TBR2/EOMES antibody (1:1000, rat monoclonal, clone Dan11mag; eBioscience at Thermo Fisher) and the KI67 antibody (1;1000, rat monoclonal, clone SolA15; eBioscience at Thermo Fisher) were conjugated with eFluor660. The primary antibodies were: Satb2 (1:1000, mouse monoclonal, ab51502, abcam), Tbr1 (1:1000, rabbit polyclonal, ab31940, abcam), GFP (1:1000, chick polyclonal, GFP-1020, aves), Brn-2 (1:1000, goat polyclonal, sc-6029, Santa Cruz), GFAP (Rabbit polyclonal, 23935-1-AP, proteintech), CTIP2 (1:1000, rat monoclonal, ab18465, abcom), ROR beta ( 1:1000, mouse monoclonal, clone N7927, perseus proteomics), Ezh2 (1:1000, mouse monoclonal, 5246S, Cell Signaling Technology), BrdU(1:1000, mouse monoclonal, clone MoBU-1, Thermo Fisher) and H3K27me3 (rabbit monoclonal,Cell Signaling Technology, #9733).

For BrdU detection, the sections were treated with 2N HCL for 1 h at 37°C after HistoVT One treatment.

#### EdU/BrdU labeling and detection

Pregnant mice were intraperitoneally injected with EdU (12.5 mg/kg body weight; Invitrogen) or BrdU(500 mg/mice; Nacalai Tesque). EdU incorporation into DNA was detected using the Click-iT™ EdU Imaging Kit (Invitrogen) according to the manufacturer’s instructions. BrdU was detected using immunostaining (see above). To trace two time points in the same mice, we injected BrdU at E12.5 and EdU at E14.5 or E15.5 and later detected each label separately (BrdU by immunostaining; EdU by click chemistry).

#### Plasmid and gRNA

The target sequences of the gRNAs used for CRISPR/Cas9-mediated knockout of Ezh2 in ferrets are listed below.

EZH2 gRNA set 1:

gRNA1:5′-GAGCTCATTGCGCGGGACCAGGG-3′

gRNA2:5′-GTGGTGGATGCAACCCGCAAGGG-3′

EZH2 gRNA set 2:

gRNA1: 5′-CTCTGCACAGACTTAGCGTTTGG-3′

gRNA2: 5′-CCATCGGACAAAGATCAAATATC-3′

Primers for gRNA synthesis were designed according to a previously described protocol (*10*). Self-amplifying PCR was performed using these primers, and the resulting amplicons were cloned into AflII-digested gRNA expression vectors modified from the Church Lab system(*11*).

All plasmids were purified using the NucleoBond Xtra Midi EF kit (Macherey–Nagel), ensuring endotoxin-free quality suitable for in vivo applications.

#### Pharmacological treatments in vitro and in vivo

For in vitro experiments, 2 × 10⁵ cells were cultured in 4-well plates in 500 μL of culture medium consisting of 20 ng/mL human basic FGF (Peprotech), 1× B27 supplement without retinoic acid (Gibco), and DMEM/F12 with GlutaMAX (Gibco). Cells were treated with DMSO (vehicle control), DMOG (150 μM or 300 μM), or α-KG (1 mM or 5 mM).

For in vivo DMOG treatment, pregnant mice were intraperitoneally injected with saline or 8 mg/mice DMOG from E11.5 to E15.5.

#### Library construction for single cell RNA-seq using ferret

Dissociation and sorting of GFP+ cells from the control and gRNA sets treated ferret cortex were performed as described above. After sorting, the cell numbers were counted using a Countess II cell counter (Invitrogen).

The collected cells were immediately loaded onto the 10X-Genomics Chromium (10X Genomics, Pleasanton, CA). Libraries for single-cell cDNA were prepared using the Chromium Next GEM Single Cell 3′ Reagent Kits v3.1 (Dual Index) platform, following the manufacturer’s protocol.

#### Single-cell RNA-seq analysis and pseudotime estimation

Single-cell RNA-seq data from ferret E32 and E40 (control and Ezh2 KO) were processed using the Seurat R package (v5.2)(*12*). Low-quality cells (fewer than 200 genes or >20% mitochondrial reads) and lowly expressed genes (in <3 cells) were excluded. The data were log-normalized and scaled. Principal component analysis (PCA) was performed using 2000 variable genes, followed by UMAP for visualization and graph-based clustering (resolution = 0.5). Differentially expressed genes (DEGs) were identified using the FindMarkers function with default parameters.

To estimate developmental progression, we calculated a temporal score (birthday score) based on the weighted expression of the curated temporal identity genes, as described above.

#### RNA-seq Library Preparation and Sequencing

RNA-seq libraries were prepared according to the manufacturer’s protocols using the following kits: NEBNext Poly(A) mRNA Magnetic Isolation Module (NEB #E7490) for mRNA enrichment, NEBNext Ultra II Directional RNA Library Prep Kit for Illumina (NEB #E7760) for strand-specific library construction, and NEBNext Multiplex Oligos for Illumina (96 Unique Dual Index Primer Pairs) (NEB #E6440) for indexing. Sequencing was performed on an Illumina NovaSeq 6000 system using the SP 100-cycle kit in the 61 bp paired-end mode or Nova-Seq Xplus in the 150-cycle paired-end mode.

#### Bulk RNA-seq processing and differential expression

Paired-end FASTQ files were quality-trimmed with Trim Galore (Phred ≥30; minimum length 50 bp; paired mode), then aligned to the mouse or ferret reference genome (GRCm39, refdata-gex-GRCm39-2024-A) using HISAT2(*13*). SAM files were converted, sorted, and indexed with samtools (*14*). Gene-level counts were obtained with featureCounts (Subread)(*15*), counting read pairs over exons and summarizing by the gene_name attribute (-p --countReadPairs -t exon -g gene_name).

Downstream analysis was performed using R with DESeq2(*16*). Raw counts and sample metadata (condition × time) were used to build a DESeq2 dataset (design = ∼ condition). Differential expression was computed with Wald tests and Benjamini–Hochberg FDR control; genes with adjusted p < 0.05 were considered significant. For visualization and unsupervised QC, we applied the DESeq2 variance-stabilizing transformation (VST; vst(blind = FALSE)) and plotted PCA from VST values. Heat maps were drawn for the top DE genes using row-wise z-scores. Volcano plots marked |log2FC| > 1 and FDR < 0.05. For day-specific contrasts, we subset samples by day and tested kg vs con; where noted, group differences in VST expression were additionally summarized as mean ± SE and compared with two-sided Welch’s t-tests (and Wilcoxon rank-sum as a robustness check), reporting BH-adjusted p-values when multiple comparisons were made.

#### Western Blot Analysis

Cells were lysed with Pierce® IP lysis buffer (Thermo Fisher Scientific), sonicated (TOMY HandySonic, 10 s, level 4), and centrifuged (13,000 × g for 10 min). Supernatants were mixed with a protease inhibitor cocktail (Nacalai Tesque) and Laemmli buffer (Sigma), boiled at 98 °C for 5 min, separated by SDS-PAGE (SuperSep, WAKO), and transferred to 0.2 μm nitrocellulose membranes (GE Healthcare). Membranes were blocked in 5% milk/TBST, incubated with primary antibodies overnight at 4 °C, followed by HRP-conjugated secondary antibodies (GE Healthcare). Signals were detected using Chemi-Lumi One Ultra (Nacalai Tesque) and imaged with an ImageQuant LAS500 system (Cytiva). Band intensity was quantified using ImageJ software. Primary antibodies: H3K27me3 (Cell Signaling Technology #9733), α-tubulin (1:1000, mouse monoclonal, DM1A, Sigma-Aldrich #T9026).

#### Human cerebral organoids culture and compound treatment

Human cerebral organoids were generated from human embryonic stem cells (hESCs; line: KhES-1 and) using the SFEBq-based 3D culture protocol described by (*17*), with minor modifications for small-molecule treatments. On day 0, hESCs were dissociated into single cells using TrypLE Express (Thermo Fisher Scientific #12605010) and seeded at 9,000 cells/well in low-adhesion 96-well V-bottom plates. Cells were cultured in 100 μL of GMEM-based medium (GMEM, Thermo Fisher Scientific #11710-035) supplemented with 20% KnockOut Serum Replacement (Thermo Fisher Scientific #10828028), 1 mM sodium pyruvate, 0.1 mM nonessential amino acids, 0.1 mM 2-mercaptoethanol, 20 μM Y-27632, 3 μM IWR-1-endo, 5 μM SB431542, and 1% penicillin-streptomycin under standard conditions (37°C, 5% CO₂). On day 3, an additional 100 μL of the same medium was added to each well. From days 6 to 18, half of the medium was exchanged every 3–4 days with fresh medium of the same composition, excluding Y-27632. From day 18, aggregates were transferred to EZ-SPHERE dishes and maintained in a neural differentiation medium consisting of DMEM/F12 supplemented with N2 supplement (100×, Thermo Fisher Scientific #17502048), Chemically Defined Lipid Concentrate (1000×, Thermo Fisher Scientific #11905-031), 1% penicillin-streptomycin, and Amphotericin B (1000×, Thermo Fisher Scientific #15290-018), under high oxygen conditions (37°C, 40% O₂, 5% CO₂).

Starting on day 34, the culture medium was changed to DMEM/F12 supplemented with N2 supplement (100×), Chemically Defined Lipid Concentrate (1000×), 10% fetal bovine serum (FBS), 5 μg/mL heparin, Amphotericin B (1000×), and 1% Matrigel (Corning #356230).

On days around 35, organoids exhibiting radial neuroepithelial organization were transferred into individual dishes and treated with one of the following compounds: DMSO (vehicle control), 2.5 mM α-Ketoglutarate (cell-permeable form), 5 mM α-Ketoglutarate,10 μM GSK343 (EZH2 inhibitor). All treatments were refreshed every three to four days. From day 50, the organoids were transferred to gas-permeable Lumox dishes (Sarstedt #94.6077.305) to enhance the oxygen exchange. Organoids were harvested on days 52 (D52) and 65 (D65) for downstream analysis.

To isolate human neural stem cells (NSCs), a previously established PAX6::Venus knock-in hESC line was used (*17*). Organoid differentiation and culture were performed following the Matrigel-free protocol described by Sakaguchi et al., 2019 (*18*). The protocol remains unchanged, except that the culture medium was not replaced with a medium containing 1% Matrigel and 10% FBS from day 34.

#### Metabolic analysis

Metabolomic analysis was performed on mouse cortical (E13) and human cortical organoid (D54) samples (n = 4 for each group). The samples were weighed before extraction. Each sample was homogenized in 1,500 μL of 50% acetonitrile aqueous solution (v/v) containing 1 μM internal standard using a tissue disruptor under cooled conditions (1,500 rpm, 120 s × two cycles). After homogenization, the samples were centrifuged at 2,300 × g for 5 min at 4°C. Supernatants were transferred to 5 kDa molecular weight cutoff centrifugal filter units (Ultrafree MC PLHCC, HMT) and filtered by centrifugation at 9,100 × g and 4°C. The filtrates were evaporated to dryness and reconstituted in Milli-Q water for CE-MS analysis. Metabolomic profiling was performed in both cation and anion modes using capillary electrophoresis–time-of-flight mass spectrometry (CE-FTMS). Cation mode: CE was performed using an Agilent CE system coupled to a Q Exactive Plus mass spectrometer. Fused silica capillaries (50 μm i.d., 80 cm) were used with a Cation Buffer Solution (HMT #H3301-1001). Electrospray ionization (ESI) was performed in the positive mode (capillary voltage: 4,000 V), and the scan range was 60–900 m/z. Anion mode: The same instrumentation was used with an Anion Buffer Solution (HMT #I3302-1023). ESI was performed in the negative mode (capillary voltage: 3,500 V), and the scan range was 70– 1, 050 m/z. Raw CE-MS data were processed using the MasterHands software developed at Keio University. Peaks with a signal-to-noise ratio ≥ 3 were extracted, and peak alignment was performed across samples based on the m/z and migration time (MT). Adduct ions (e.g., Na⁺ and K⁺) and fragment ions were removed unless they were compound-specific. The relative peak areas were normalized using an internal standard and sample weight. For metabolite identification, the peak m/z and MT values were matched against the HMT compound library within tolerances of ±0.5 min for MT and ±5 ppm for m/z. Quantification was performed using one-point calibration curves with internal standard normalization. A total of 353 compounds were targeted, and 105 metabolites (52 cationic and 53 anionic) were successfully quantified.

#### Statistics

For each experiment, we first assessed the normality of distribution using the Shapiro–Wilk test. If all groups followed a normal distribution (*p* > 0.05), we tested for homogeneity of variance using Levene’s test. When both assumptions were satisfied, we performed one-way analysis of variance (ANOVA), followed by Tukey’s Honest Significant Difference (TukeyHSD) post-hoc test to evaluate the pairwise group differences.

If the data did not meet the assumption of normality, we used the non-parametric Kruskal–Wallis test, followed by Dunn’s test with Holm correction for multiple comparisons.

To assess the effects of different drugs on organoid development, we compared each treatment group with the control group using the Wilcoxon rank-sum test (also known as the Mann–Whitney U test) because the data did not satisfy the assumptions of normality.

To correct for multiple comparisons, the FDR correction was applied using the Benjamini–Hochberg procedure. Adjusted *p*-values less than 0.05 were considered statistically significant. All statistical analyses were conducted in R.

## Supplementary Text

### Mathematical Modelling

#### GRN Model

To investigate how developmental tempo is modulated across species, we abstracted the experimentally observed temporal cascade into a simplified gene regulatory network (GRN). Specifically, we replaced the three representative temporal fate genes Foxp1 (deep-layer neurons), Brn2 (upper-layer neurons), and Zbtb20 (glial cells) with hypothetical variables F, B, and Z to focus on their regulatory relationships rather than their molecular details. The GRN was designed to reproduce the sequential activation pattern observed in vivo (Fig. 1A to C): F is expressed first and later repressed by B; once F is downregulated, Z is upregulated, and Z in turn represses B, completing the cycle (Fig. 1D).

We modeled this GRN as a set of ordinary differential equations, resulting in three mRNA equations and three protein equations:

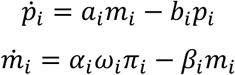

where *p_i_* is the protein concentration, and *m_i_* is the mRNA concentration of gene 𝑖 = 𝐹, 𝐵, 𝑍. The parameters *a_i_* and 𝛼*_i_*, are the translation and transcription rates respectively, while 𝑏*_i_* and 𝛽*_i_* are the degradation rates of protein and mRNA respectively.

Repression is introduced via 𝜔*_i_*, the probability that the gene is transcriptionally available, regulated by the presence of proteins from each gene:

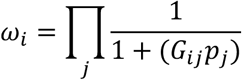

The parameter *G_ij_* represents the strength of the repression of gene 𝑗 on gene 𝑖. To introduce the role of methylation, we also include 𝜋_𝑖_, the probability that the regulatory region of gene 𝑖 is accessible to the transcriptional machinery (i.e not methylated), which, combined with 𝜔_𝑖_, determines the speed of transcription.

#### Reduction and normalisation of equations

Now we assume that mRNA dynamics are faster than protein dynamics. Following quasi steady state reduction, we find that when 𝑚_!_ is at equilibrium,

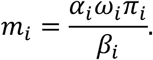

Plugging this into the original ODE, we find that

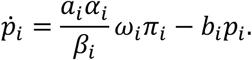

We normalise by defining the maximum of *p_i_* as 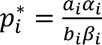, we peform a change of variables with 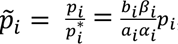, giving us

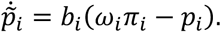

The degradation rate 𝑏*_i_*. appears as a prefactor controlling the overall speed of the system. From now on, we drop the tilde. Multiplying all 𝑏*_i_* by a constant factor proportionally scales the tempo, e.g halving all protein degradation rates would make the system run twice as slow.

Guided by experimental results showing that these temporal genes gradually lose methylation over time (Fig 2. C to E), we model the methylation function as a dynamic variable:

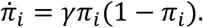

We will call this form of 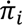 as the ‘autonomous’ model, since its dynamics are only affected by its own value, i.e protein concentrations have no effect.

Throughout the paper, when calculate the ‘protein half-life’ via log (2)⁄𝑏, and ‘methylation half-life’ as log (2)⁄𝛾. Note that this is only technically the methylation half-life at 𝜋 = 0, i.e at the beginning of the simulation, giving us an approximate measure of the timescale of demethylation.

#### Computational simulations (Fig.1D-I)

The simulations of the mathematical model were carried out in Python, with the integration performed using scipy’s ‘solve_ivp’ function. The initial conditions are set as (𝑝_1_, 𝑝_2_, 𝑝_3_, 𝜋_1_, 𝜋_2_, 𝜋_3_) = (0.5,0,0,1,0.1,0.1).

The parameter set used For Fig.1E and F is shown in the following table (for Fig.1F, all degradation rates are scaled uniformly):

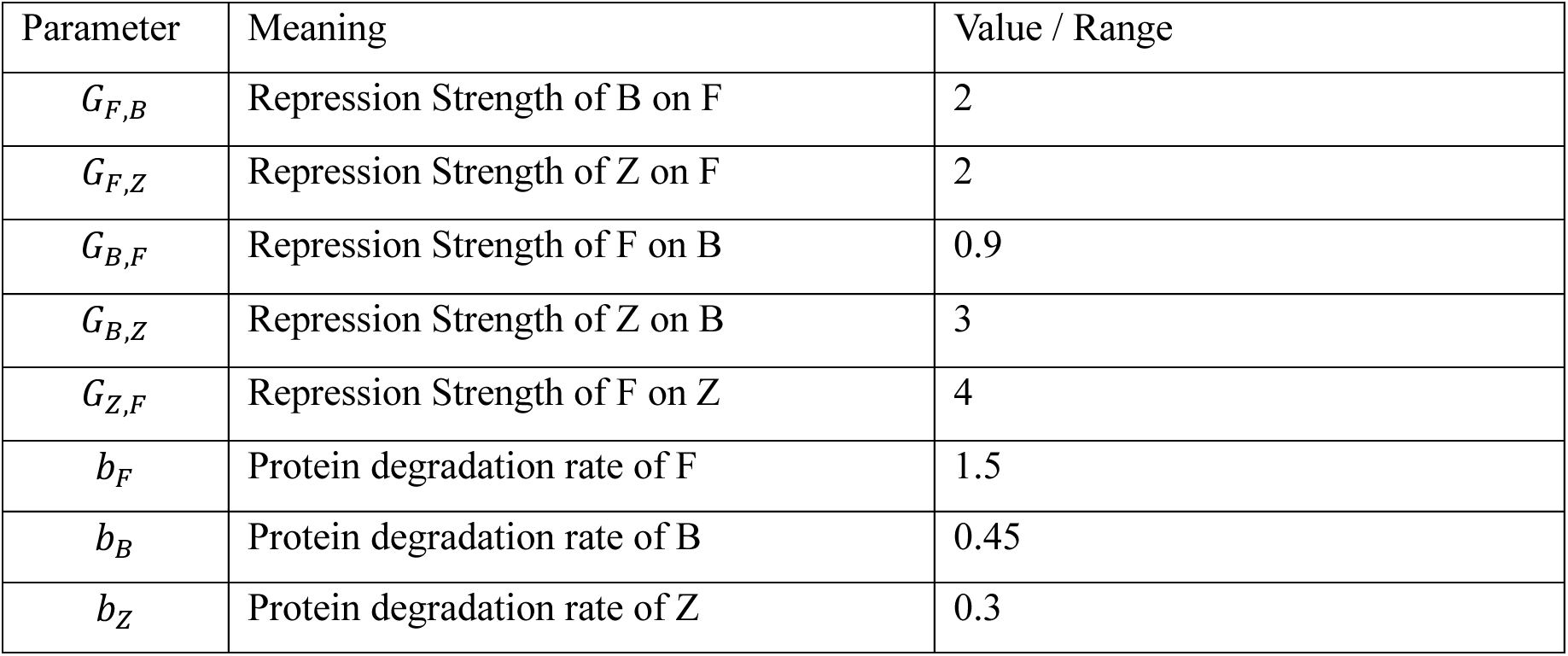

The parameter set used for Fig.1H is shown in the following table.

| Parameter | Meaning | Value / Range |
| --- | --- | --- |
| $G_{F,B}$ | Repression Strength of B on F | 1 |
| $G_{F,Z}$ | Repression Strength of Z on F | 2 |
| $G_{B,F}$ | Repression Strength of F on B | 1 |
| $G_{B,Z}$ | Repression Strength of Z on B | 1 |
| $G_{Z,F}$ | Repression Strength of F on Z | 2 |
| $b_B/b_F$ | Relative protein degradation rate of B | 1 |
| $b_Z/b_F$ | Relative protein degradation rate of Z | 1 |
| $b_F$ | Protein degradation rate of F | 0.001–0.058 hr<sup>-1</sup> (half-life ≈ 0.5–3 days) |
| $\gamma$ | Methylation rate | 0.001–0.116 hr<sup>-1</sup> (half-life ≈ 0.25–20 days) |

To investigate how degradation and methylation rates contribute to developmental tempo (Fig.1I), we systematically explored a range of parameter combinations using a parameter search. We used Latin hypercube sampling (LHS) to sample the high-dimensional parameter space. We ran 3000 simulations, with the ranges used for Fig.1I shown in the following table:

| Parameter | Meaning | Value / Range |
| --- | --- | --- |
| $G_{F,B}$ | Repression Strength of B on F | 0.1 - 10 |
| $G_{F,Z}$ | Repression Strength of Z on F | 0.1 - 10 |
| $G_{B,F}$ | Repression Strength of F on B | 0.1 - 10 |
| $G_{B,Z}$ | Repression Strength of Z on B | 0.1 - 10 |
| $G_{Z,F}$ | Repression Strength of F on Z | 0.1 - 10 |
| $b_B/b_F$ | Relative protein degradation rate of B | 0.3 - 3 |
| $b_Z/b_F$ | Relative protein degradation rate of Z | 0.3 - 3 |
| $b_F$ | Protein degradation rate of F | 0.14 hr<sup>-1</sup> (half-life ≈ 2–20 days) |
| $\gamma$ | Methylation rate | 0.002 - 0.083 hr<sup>-1</sup> (half-life ≈ 0.5–20.5 days) |

A representative subset of these simulations is shown in the swarm plot in Fig.1I.

#### Filtering simulations for biological relevance and quantifying tempo

Before plotting each simulation, we checked that it fit the criterion of the peaks of the genes in the correct order: F, then B, then Z. Only simulations that fit this criterion were plotted. For genes F and B, which exhibited well-defined transient maxima, the peak was identified as the time point at which the expression reached its maximum value.

For gene Z, which typically monotonically increases, identifying the time of the true maximum is less informative. In many cases, the maximum occurs at or near the final simulation time point, making it a poor indicator of the tempo of expression. Instead, we defined the peak for Z as the first time point at which the expression exceeded 90% of its maximum value.

The tempo is quantified via the ‘genetic program duration’, which is the time taken to reach the Z peak (i.e., it measures how slow the program is).

---

**Fig. S1.**
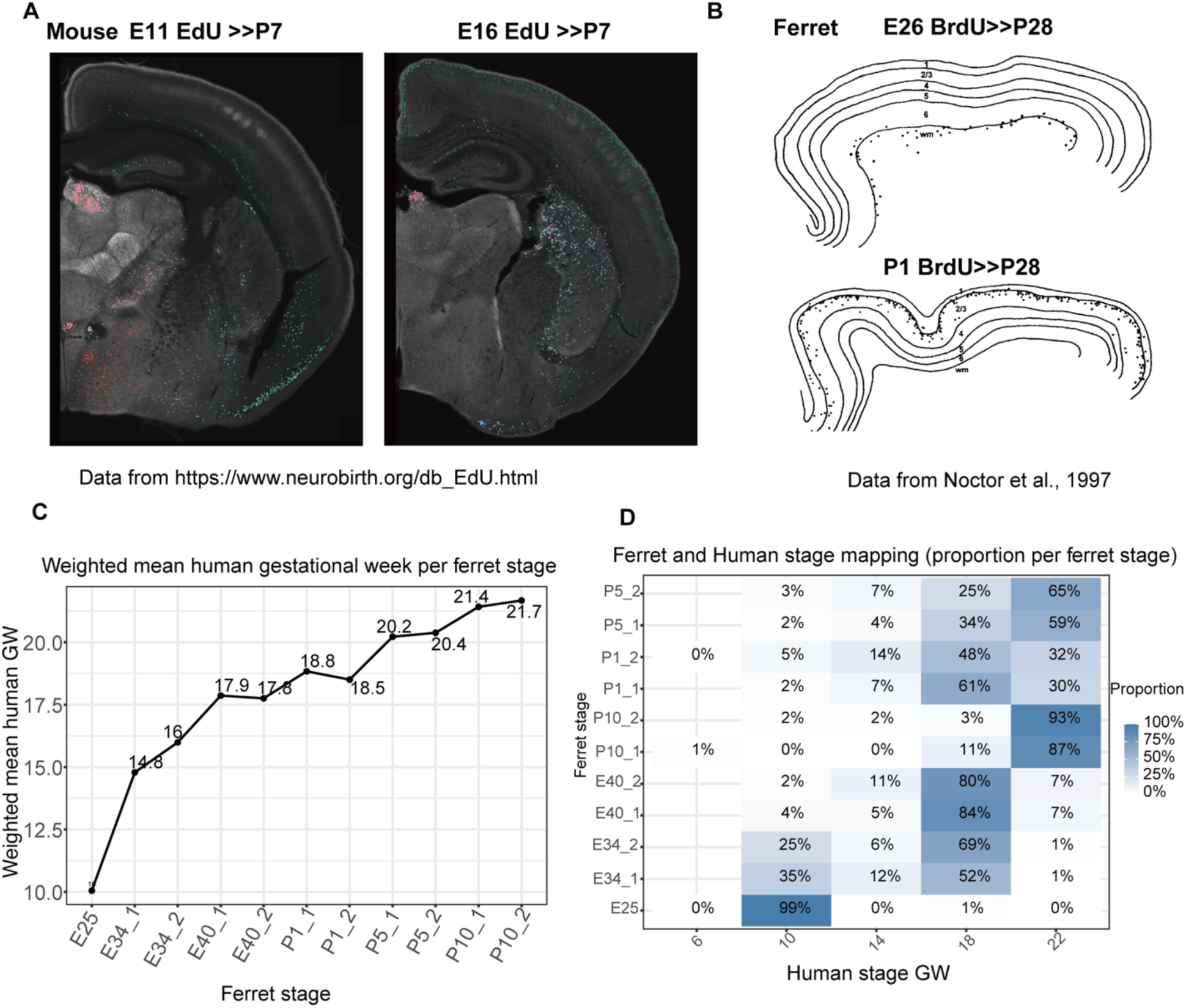
Cross-species alignment of cortical neurogenesis. (**A**) Mouse EdU birth-date tracing: labels at E11 (left) and E16 (right) with analysis at P7, showing the deep-layer onset and the end of upper-layer production of neurogenesis. Data were obtained from the NeuroBirth EdU database (https://www.neurobirth.org). (**B**) Ferret BrdU birth-date tracing: pulses at E26 (top) or P1 (bottom) with analysis at P28, illustrating the deep-layer onset and the end of upper-layer production of neurogenesis, similar to that observed in mice. Schematic adapted from (*4*). (**C**) Weighted mean human gestational week (GW) predicted for each ferret stage (from E25 to P10) using Seurat anchor-based label transfer from human scRNA-seq references (*19*); numbers denote the weighted mean GW. (**D**) Heatmap of the proportion of human GW assignments for each ferret stage (rows) derived from the same label-transfer predictions as in C. The concentration of mappings within GW10–18 provides a rationale for using this window to compare ferret and human neurogenesis.

**Fig. S2.**
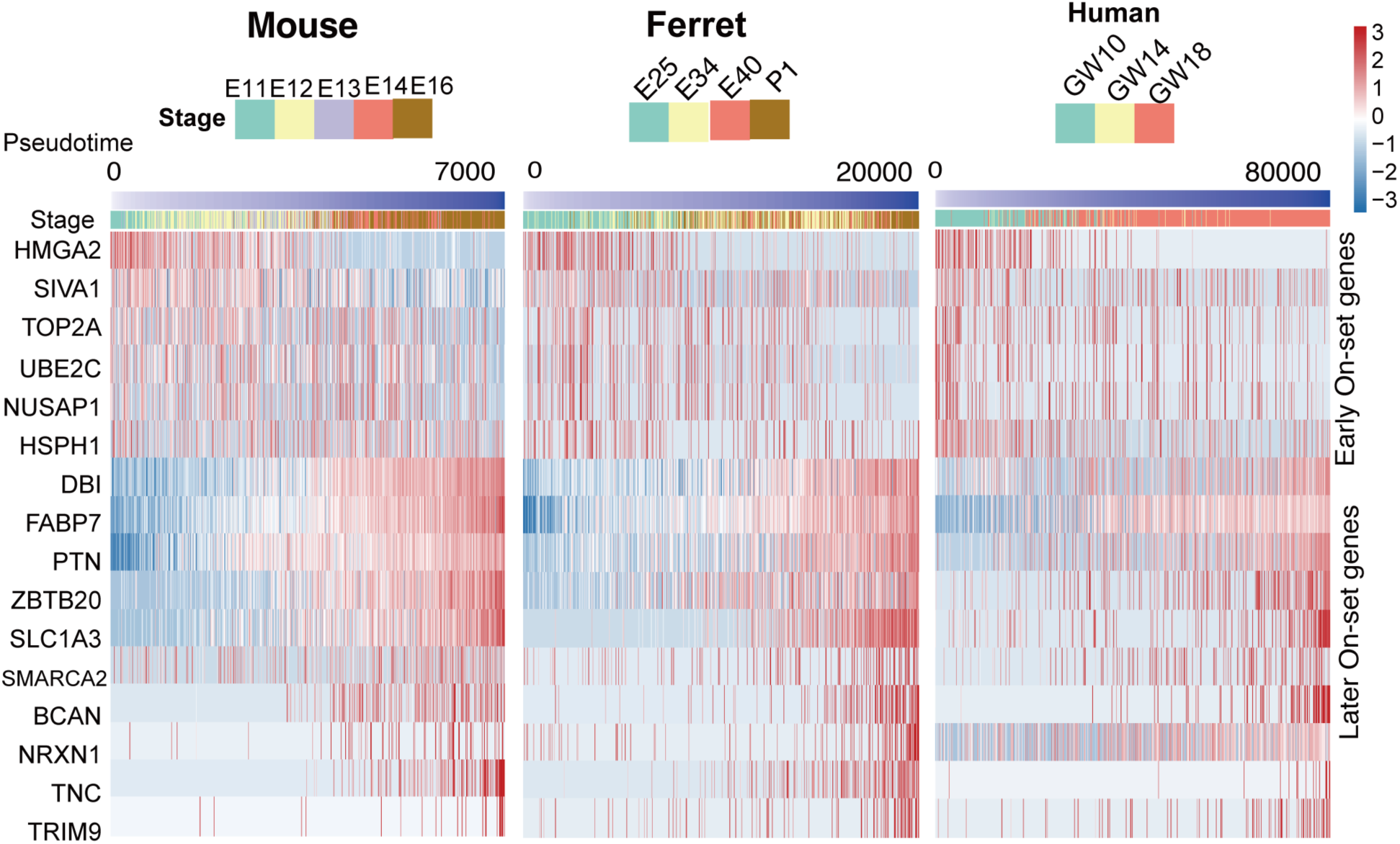
Cross-species alignment of cortical neurogenesis. Heatmap of the expression patterns of several temporal genes across three species: mice, ferrets, and humans. Gene expression levels were normalized and aligned to the developmental pseudotime.

**Fig. S3.**
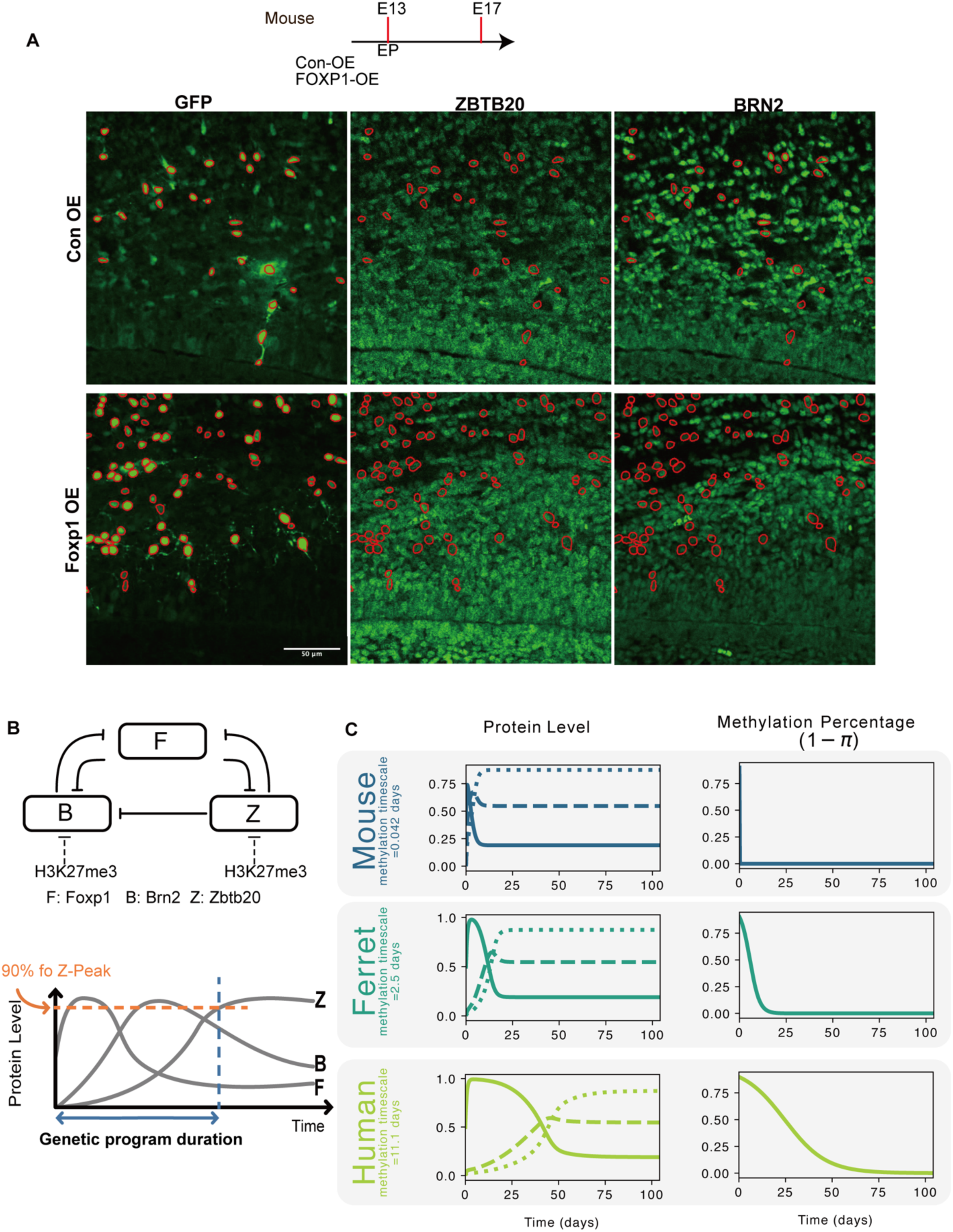
FOXP1 overexpression suppresses late temporal gene expression in the embryonic mouse cortex and a mathematical model illustration. **(A)** Experimental timeline (top): *in utero* electroporation (IUE) at E13 and analysis at E17. Representative coronal sections from control overexpression (Con-OE) and FOXP1 overexpression (FOXP1-OE) brains. GFP marks electroporated cells (left). Immunostaining for ZBTB20 (middle) and BRN2 (right); red circles indicate GFP⁺ cells co-expressing the indicated markers. Compared with Con-OE, FOXP1-OE showed a decreased fraction of GFP ^+^ cells expressing the late temporal markers ZBTB20 and BRN2. (Scale bar as indicated.) **(B)** Mathematical model and definition of the duration of the genetic program. The duration is determined from the initiation of the simulation (time 0) to the point at which the Z-peak attains 90% of its maximum value (indicated by the red line) during the simulation period.**(C)** Model outputs used in the study: example trajectories of protein levels (left) and methylation fraction (1–π) (right) under different parameter sets corresponding to mice, ferrets, and humans. Slower demethylation yields delayed induction and prolonged persistence of late temporal proteins, providing a conceptual framework for interpreting shifts in marker onset.

**Fig. S4.**
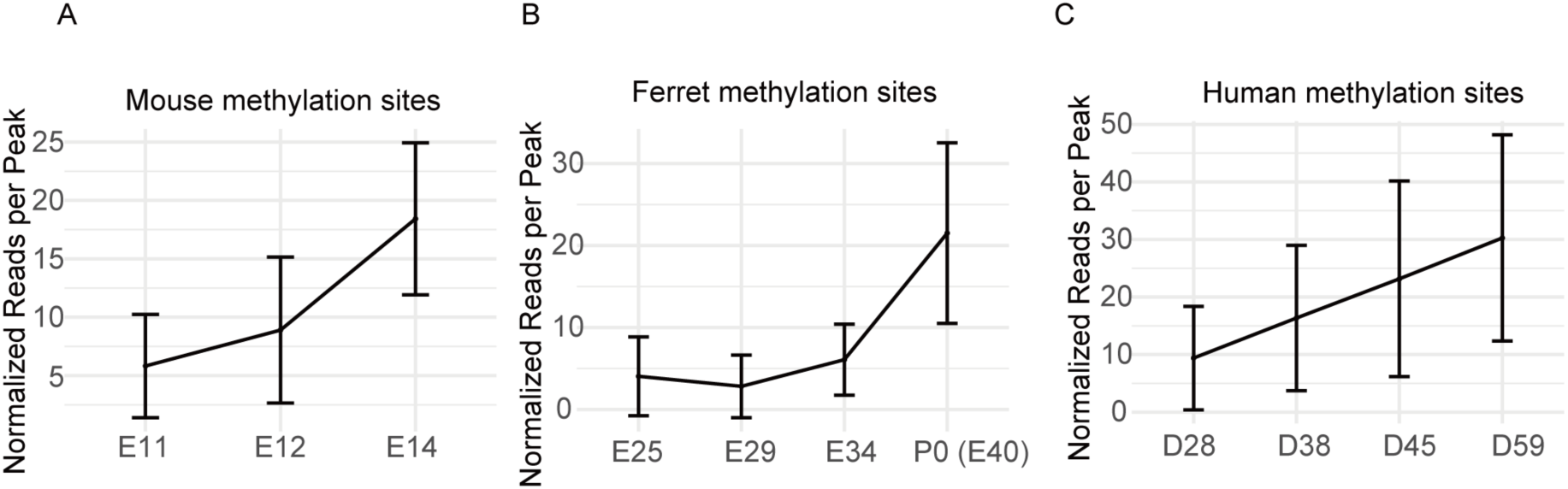
Global separation of NSC epigenomes and dynamics at methylation sites. **(A to C**) Mean normalized reads per H3K27me3 peak (DiffBind; library-normalized reads/peak) at methylation sites over developmental time in mice (A), ferrets (B), and humans (C). Points show the mean across all sites and replicates; error bars denote ±SD across peaks.

**Fig. S5.**
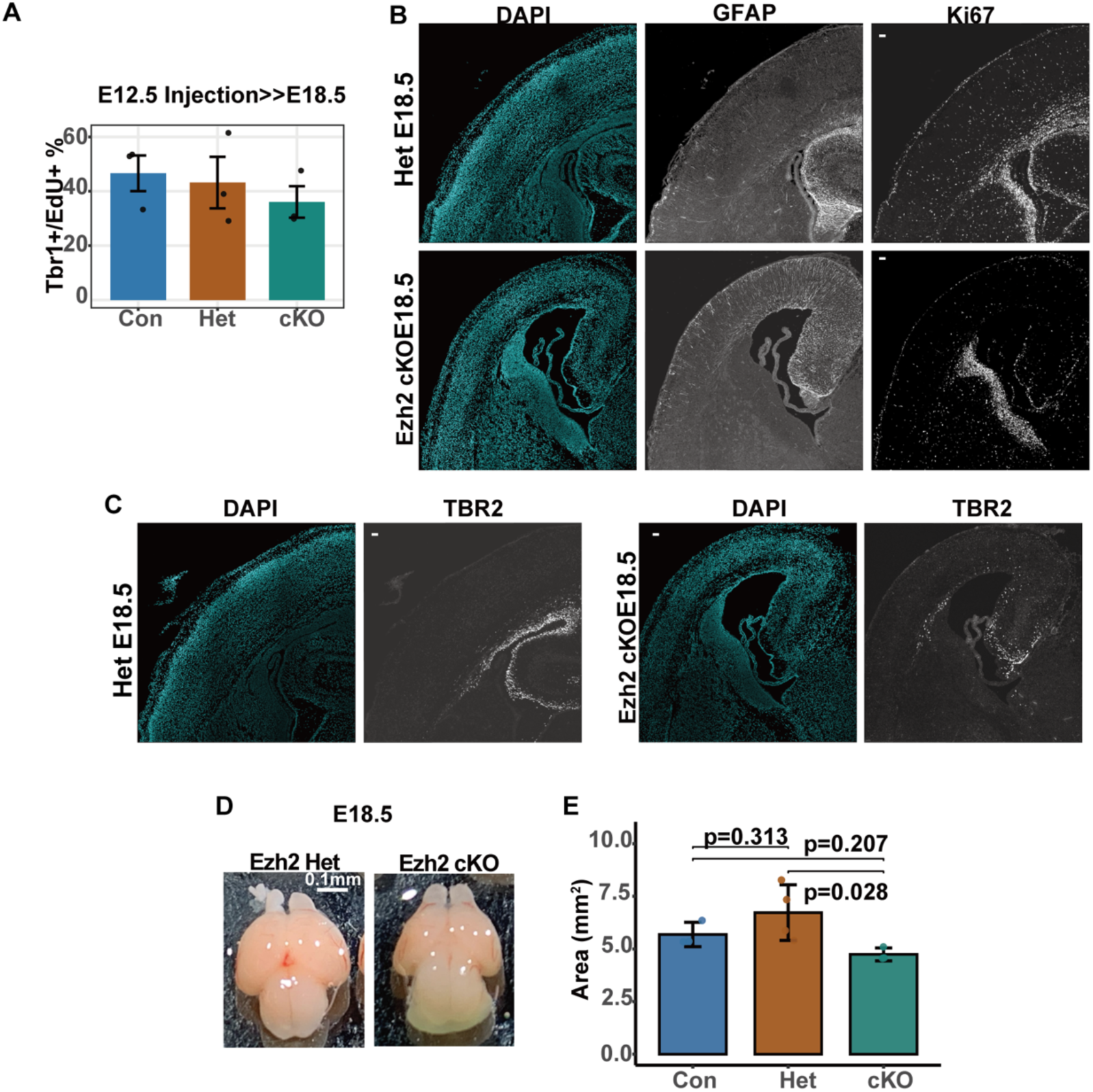
Loss of Ezh2 scales neurogenesis and gliogenesis in the embryonic mouse cortex. (**A**) Percentage of TBR1^+^EdU^+^ neurons at E18.5 in the littermates of the three genotypes after EdU injection at E12.5. The bars indicate the mean ± SD. Con: n=4; Het: n=3; cKO: n=3. (**B and C)** Representative coronal sections showing staining for GFAP/KI67 (A) and TBR2 (B) in *Ezh2* heterozygous **a**nd cKO mice at E18.5. (**D and E**) Representative dorsal views of whole brains from littermates used in this study (D) and measurements of the cortical area (mm^2^) of three different genotypes in littermates (E). Con: n=4; Het: n=4; cKO: n=3. The bars indicate the mean ± SD. P-values were calculated using the Kruskal–Wallis tests with Dunn’s test with Holm correction.

**Figure S6.**
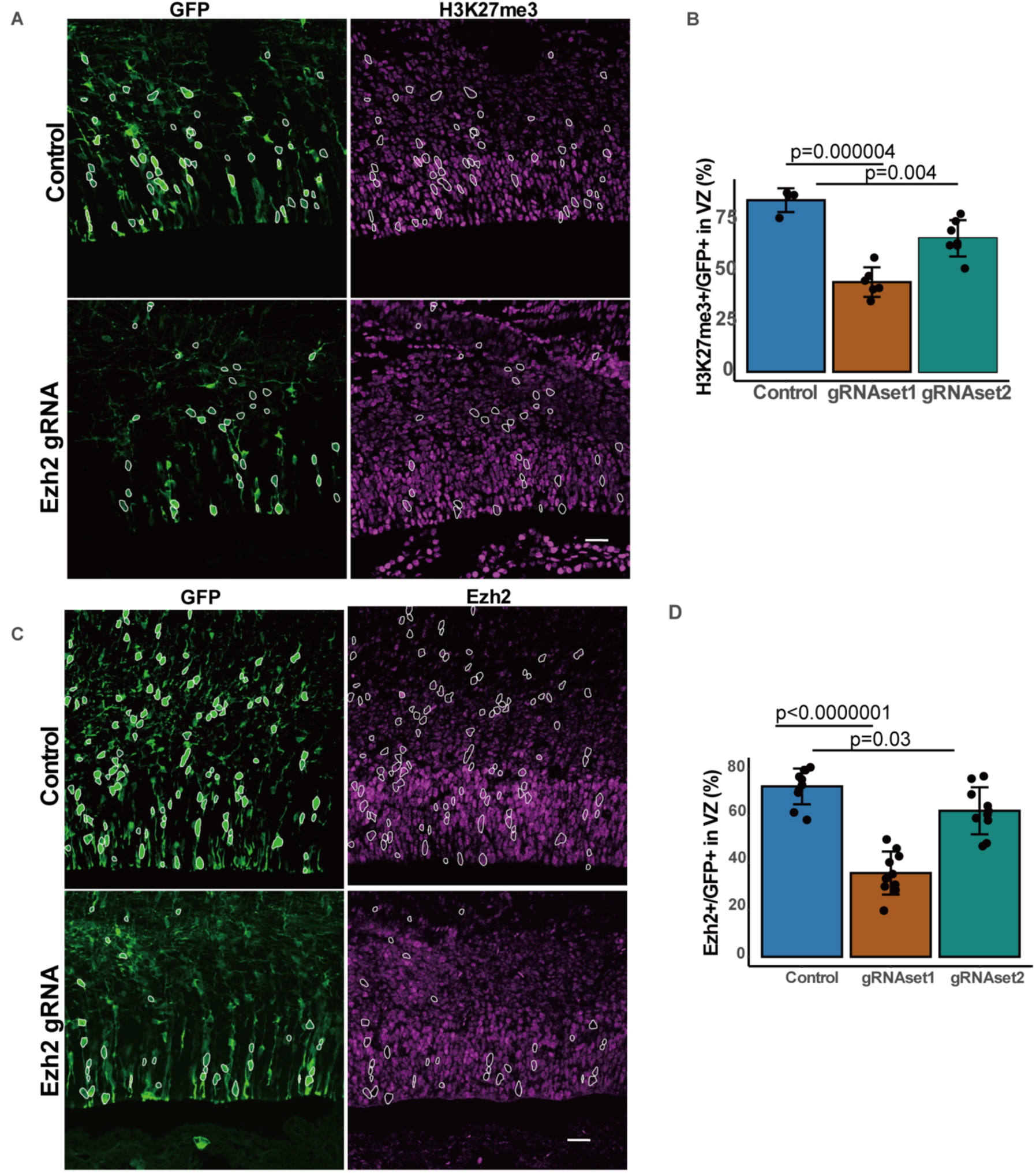
Ezh2 knockout efficiently reduced the number of H3K27me3^+^cells in the ferret cortices. Immunostaining and quantification show an efficient depletion of H3K27me3 levels **(A and B)** and Ezh2 expression **(C and D)** in the ventricular zone at E40 following the introduction of Ezh2 gRNA and CRISPR/Cas9 sets via *in utero* electroporation. P-values were calculated using ANOVA tests with Tukey’s HSD correction. Bars show mean ± SD; points represent sections from two different biological samples.

**Figure S7.**
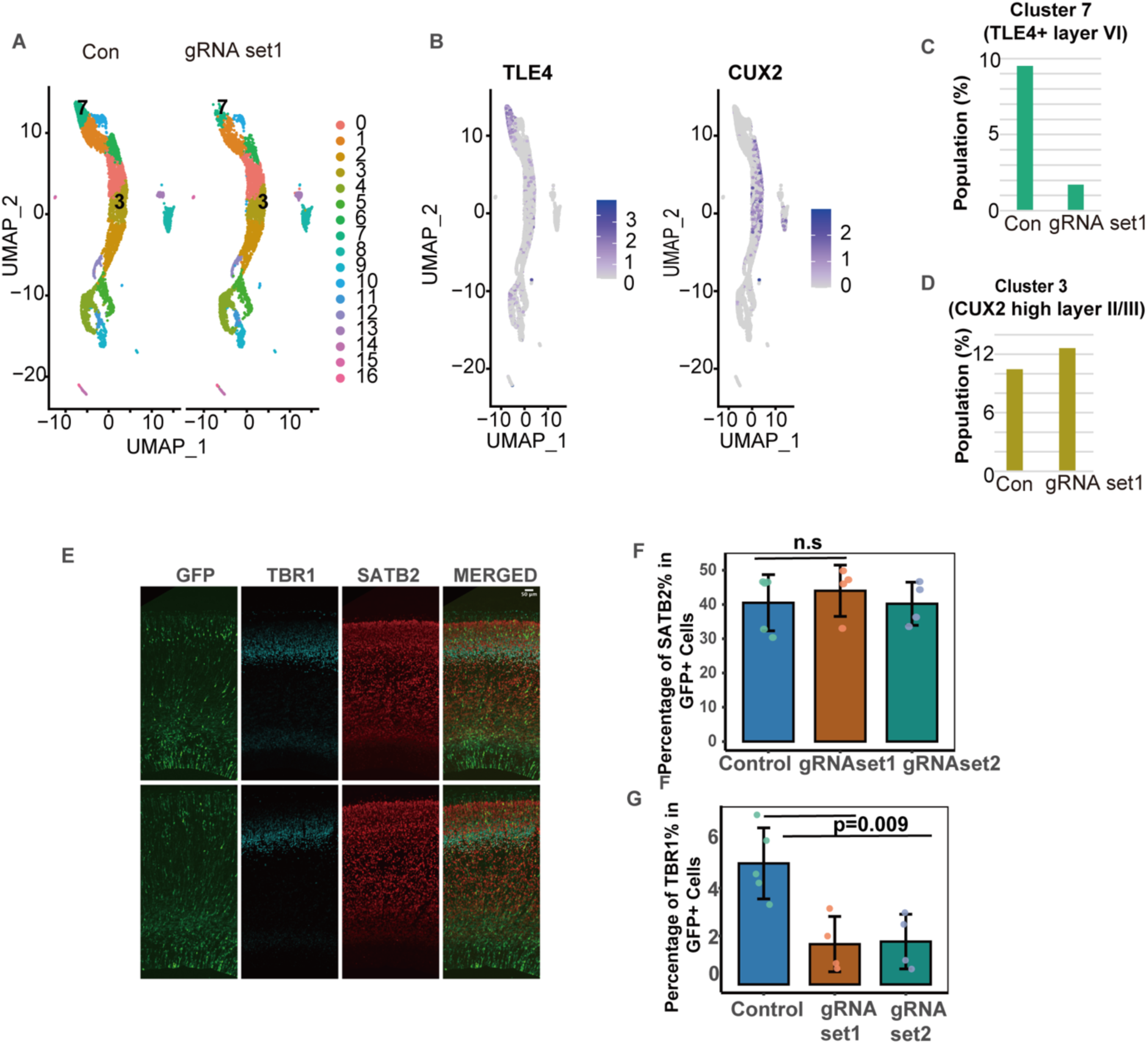
Knockout of Ezh2 affects temporal gene expression in the ferret cortex. **(A)** UMAP projection of single-cell RNA-seq data from ferret cortices at E34 following *in utero* electroporation (IUE) at E30 with GFP and no gRNA (CON) or Ezh2-targeting gRNA sets (gRNA set1). (**B)** Normalized expression of TLE4 (layer VI) and CUX2 (layer II/III) across UMAP. **(C and D)** Fraction of cells assigned to the TLE4⁺ (cluster 7) and CUX2-high (cluster 3) clusters under each condition. **(E to G)** Immunofluorescence and quantification of E40 cortices electroporated at E32 ferrets with the indicated gRNA sets showed changes in SATB2⁺ and TBR1⁺ cells in the GFP⁺ electroporated cells. P-values were calculated using ANOVA tests with Tukey’s HSD correction. Bars show mean ± SD. Con: n=5; gRNA set1: n=4; gRNA set2: n=4.

**Figure S8.**
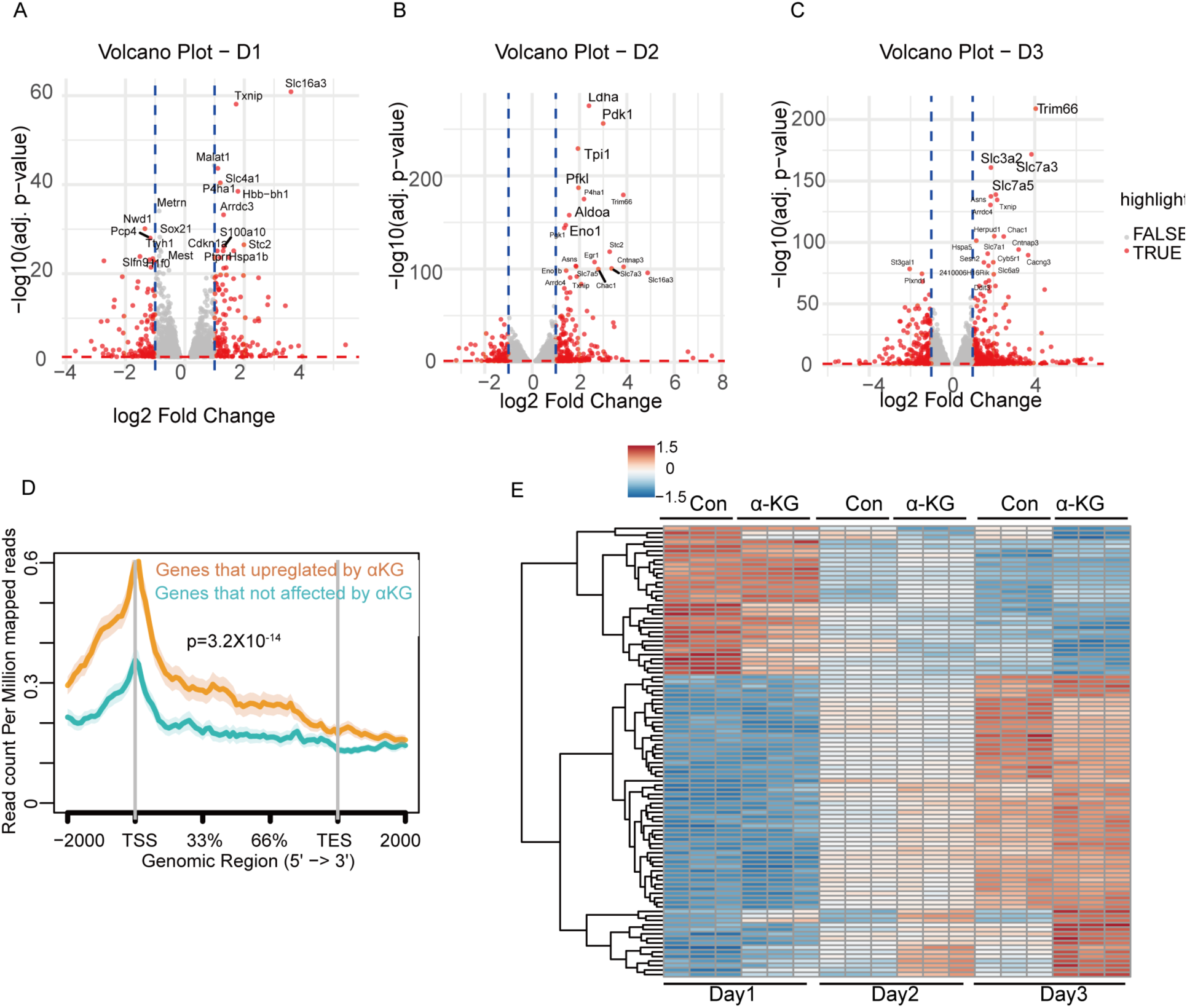
Transcriptomic changes in response to α-ketoglutarate (α-KG) across days in the mouse cortical cell cultures. **(A–C)** Volcano plots comparing α-KG (5 mM) versus control on **days 1–3**. The x-axis indicates log₂ fold change; the y-axis indicates −log₁₀(adjusted *p-value*). Red points denote significantly differentially expressed genes (DEGs; FDR < 0.05; |log₂FC| ≥ 1; vertical dashed lines). Labeled genes highlight recurrent αKG-responsive transcripts, including glycolytic (*Ldha, Pfkl, Tpi1, Eno1/2*) and transporter genes (*Slc2a1/2a3*). **(D)** Metagene profiles of H3K27me3 (from E12 NSCs) across gene bodies for genes unaffected by α-KG (top) versus genes upregulated by α-KG (bottom); shaded bands indicate ±SD. The Wilcoxon signed-rank test was used to evaluate the differences. **(E)** Heatmap of variance-stabilized expression (row-scaled z-scores) for representative α-KG-responsive genes across D1-D3. Rows were clustered, and columns were ordered by day (bottom) and treatment (top).

**Figure S9.**
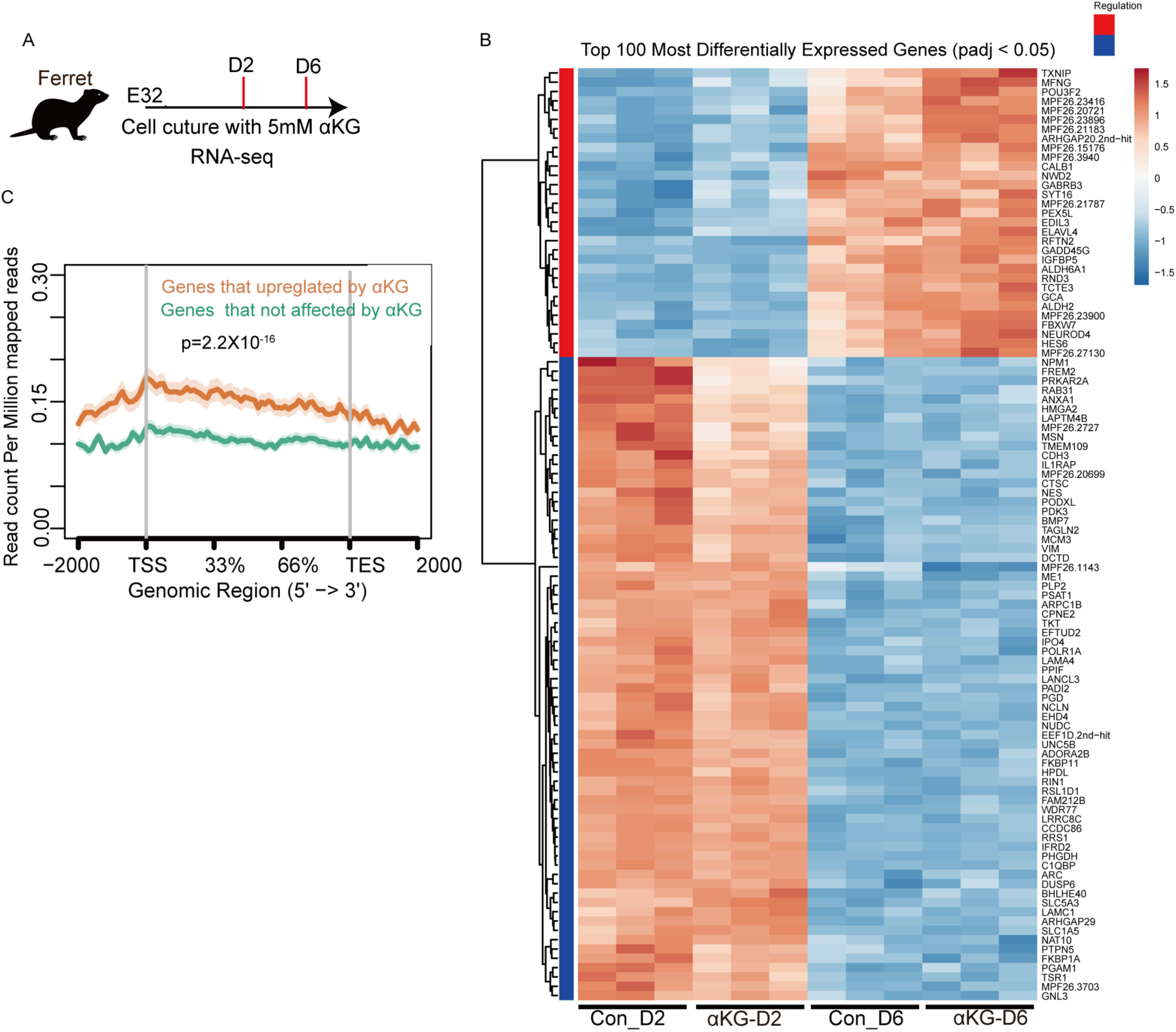
α-KG accelerates temporal gene programs and reduces H3K27me3 in ferret cortical cultures. **(A)** Experimental scheme of the study. Ferret E32 cortical cells were cultured **±**5 mM α-KG and harvested for RNA-seq at Day 2 (D2) and Day 6 (D6)**. (B)** Heatmap of the top 100 differentially expressed genes (DEGs, padj < 0.05) across the four conditions. Values are variance-stabilized and row-scaled (z-scores); the rows are clustered. Red and blue side bars indicate genes upregulated and downregulated by α-KG, respectively. **(C)** Metagene profile of H3K27me3 ChIP-seq signal across gene bodies for genes upregulated by α-KG (orange) versus those not affected (teal); shaded bands denote±SD. The Wilcoxon signed-rank test was used to evaluate the differences.

**Figure S10.**
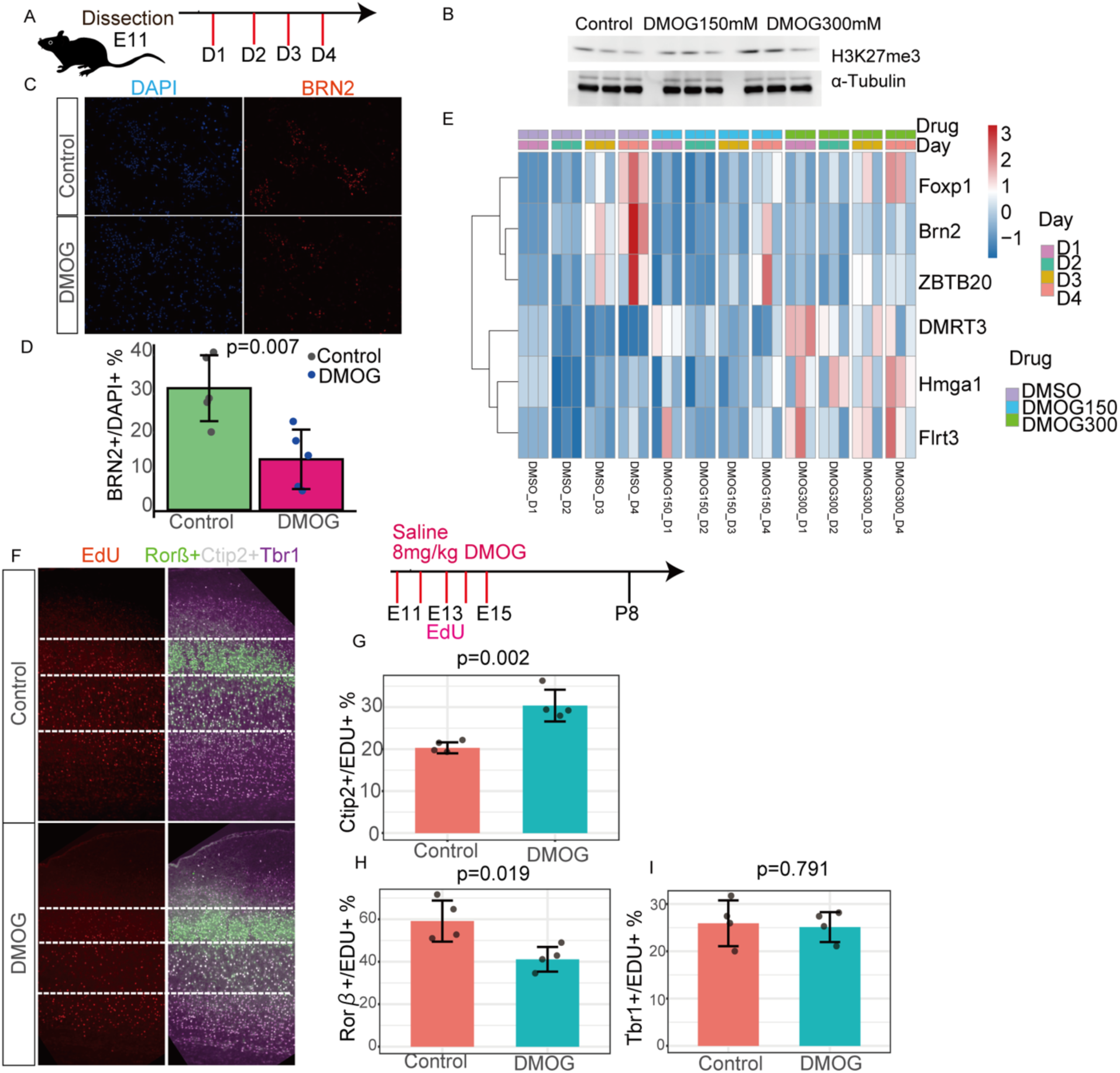
DMOG suppresses the temporal progression of cortical development both *in vitro* and *in vivo*. **(A)** Timeline of *in vitro* study. E11 mouse cortical cells were cultured with DMOG (150 or 300 µM) or DMSO and sampled on days 1–4 (D1–D4**). (B)** Western blot of H3K27me3 (α-tubulin loading control) at D2 shows an increase with DMOG treatment. **(C)** Heatmap of QPCR for representative temporal genes across D1–D4 under DMSO, DMOG150, and DMOG300 (three replicates per time point). **(D)** Immunostaining of mouse cortical cells for BRN2 and DAPI after treatment with DMSO and 300 mM DMOG for three days from E12. **(E)** Quantification of BRN2⁺/DAPI⁺ (%): bars, mean ± SD; dots, biological replicates; two-sided *p* value indicated. **(F)** Pregnant mice received DMOG (8 mg/mice) from E11 to E15 every day; EdU was administered at E13, and offspring were analyzed at P8. Representative cortical sections showing EdU (left channel) and ROR β ^+^ CTIP2 ^+^ TBR1^+^ (right), with cortical layers indicated by dashed lines. **(G–I)** Quantification of EdU-labeled neuron fate at P8: CTIP2⁺/EdU⁺ (%) increased with DMOG (G), ROR β ⁺/EdU⁺ (%) decreased (H), and TBR1⁺/EdU⁺ (%) is unchanged (I). Bars, mean ± SD; dots, biological replicates; t-test was performed to evaluate the differences.

**Fig. S11.**
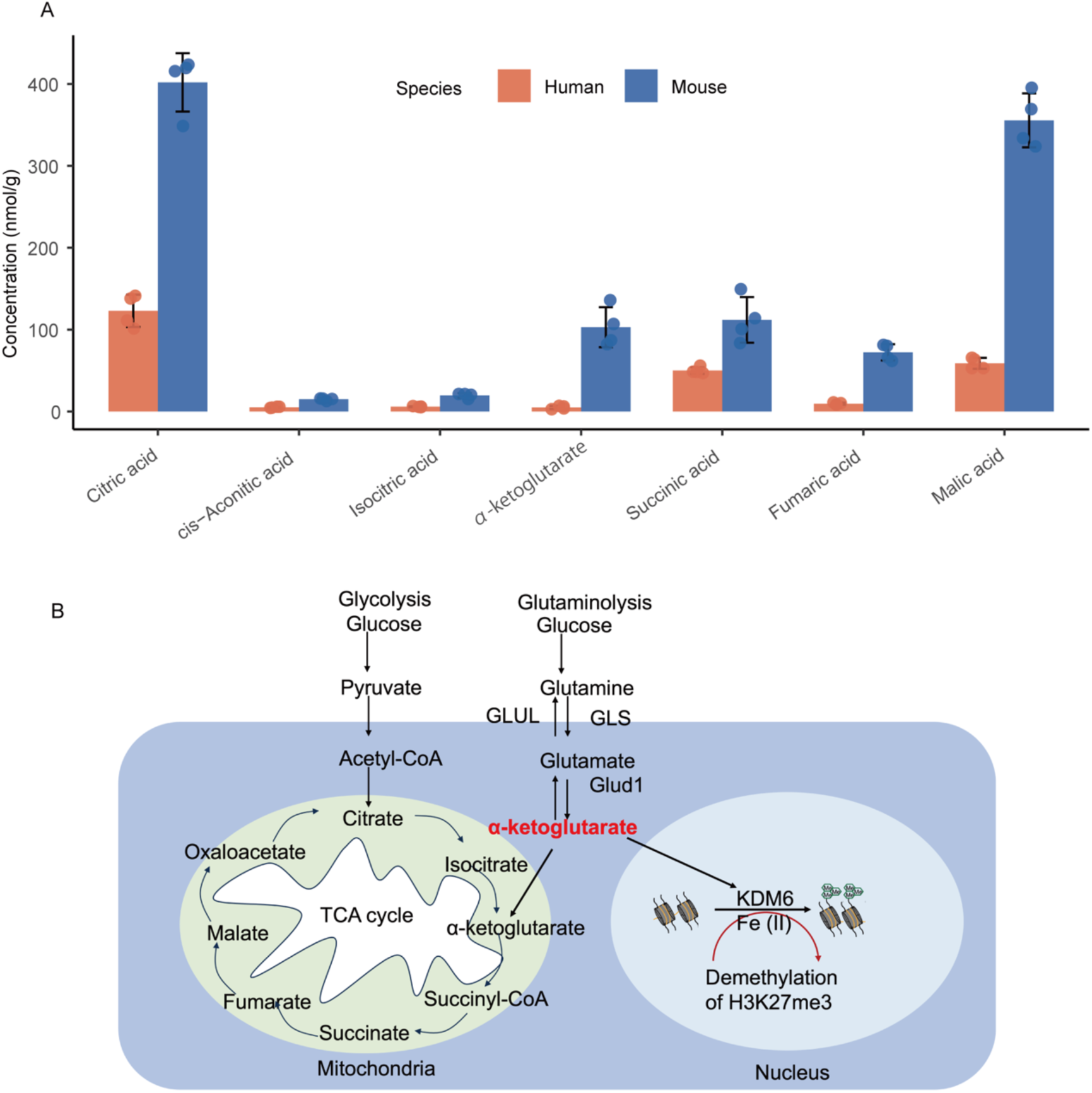
Comparing of metabolites involved in the TCA cycle between humans and mice and the α-KG–epigenetic link. **(A)** Quantification of tricarboxylic acid (TCA) intermediates in embryonic mouse cortex (E13) and human cortical organoids (D54). Bars show mean concentration (nmol g⁻¹); error bars show SD; dots represent individual biological replicates (n = 4 per species). **(B)** Schematic representation of the pathways generating α-KG, which acts as a cofactor for Fe(II)/α-KG–dependent histone demethylases of the KDM6 family (UTX/KDM6A; JMJD3/KDM6B), promoting H3K27me3 demethylation.

**Fig. S12.**
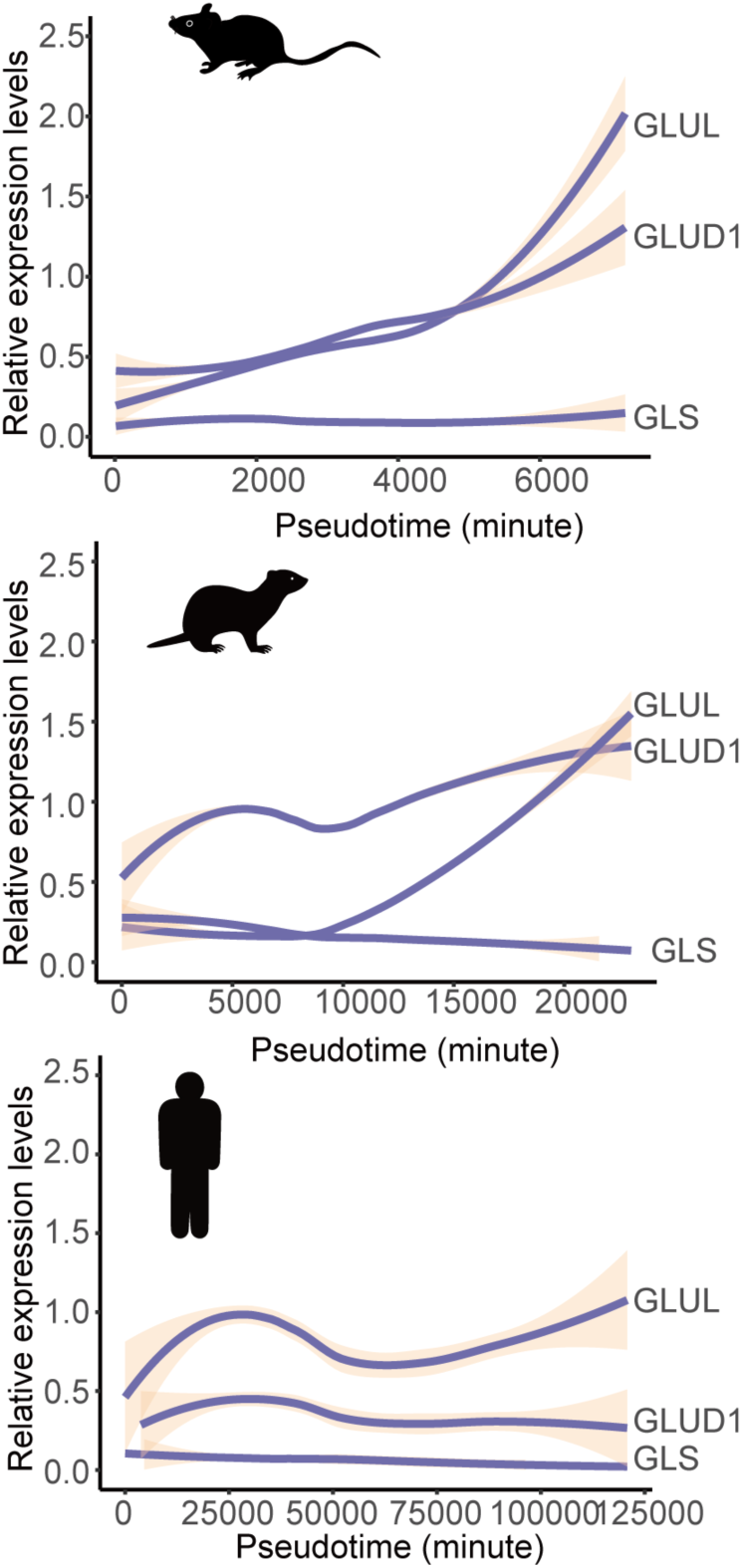
Cross-species pseudotime dynamics of glutamine metabolism genes in NSCs. Pseudotime plots showing the expression of Glud1 (glutamate dehydrogenase 1), Glul (glutamine synthetase), and Gls (glutaminase) in NSCs from mice, ferrets, and humans. Gene expression values (y-axis) were normalized within species and aligned along the developmental pseudotime.

**(separate file)**

**Table S1. List of late-onset genes in the NSCs of ferrets and humans.**

**Table S2. The number of cells in each group of control and EZH2 KO ferrets.**

**Table S3. Normalized counts for differentially expressed genes across time points (D2, D6) and α-KG treatment of ferret cortical cells.**

**Table S4. Normalized counts for top 1000 differentially expressed genes across time points (D1 to D3) and α-KG treatment of mouse cortical cells.**

**Table S5. Concentration of targeted metabolomics of in human organoid and mouse cortical samples.**

